# FBM: Freestanding bilayer microscope for single-molecule imaging of membrane proteins

**DOI:** 10.1101/2024.01.26.577465

**Authors:** Gonzalo Pérez-Mitta, Yeliz Sezgin, Weiwei Wang, Roderick MacKinnon

## Abstract

Integral membrane proteins (IMPs) constitute a large fraction of organismal proteomes, playing fundamental roles in physiology and disease. Despite their importance, the mechanisms underlying dynamic features of IMPs, such as anomalous diffusion, protein-protein interactions, and protein clustering, remain largely unknown due to the high complexity of cell membrane environments. Available methods for *in vitro* studies are insufficient to study IMP dynamics systematically. This publication introduces the Freestanding-Bilayer Microscope (FBM), which combines the advantages of freestanding bilayers with single-particle tracking. The FBM, based on planar lipid bilayers, enables the study of IMP dynamics with single-molecule resolution and unconstrained diffusion. This paper benchmarks the FBM against total internal reflection fluorescence (TIRF) imaging on supported bilayers and is used here to estimate ion channel open probability and to examine the diffusion behavior of an ion channel in phase-separated bilayers. The FBM emerges as a powerful tool to examine membrane protein/lipid organization and dynamics to understand cell membrane processes.

## Main

Integral membrane proteins (IMPs) encode 20-30% of the proteomes of organisms across all domains of life^1,2^. Receptors, transporters, and channels compose most IMPs in humans and are about 60% of all therapeutic drug targets, exemplifying the importance of these proteins in physiology and disease^3^. However, many dynamic features of IMPs remain enigmatic and controversial. For example, IMPs in the plasma membrane switch stochastically between diverse motions when observed under a microscope; the same protein can transition from a random walk to a linear motion or a complete stop^4,5^. Simultaneously, IMPs can interact with themselves and other proteins and adopt non-homogeneous distributions across membranes^6,7^. The origin of such features needs to be addressed to develop good dynamical models of cell membranes.

Explaining these phenomena requires curtailing some of the intrinsic complexities of the cellular environment. This can be achieved with experimental setups that allow the study of IMPs in membranes of controlled composition with single-particle tracking (SPT)^2,8,9^. SPT permits the study of the diverse aspects of IMP dynamics, such as anomalous diffusion, dynamic heterogeneity, and protein-protein interactions, by obtaining the full information contained in particle positions over the course of a time-lapse experiment^10–12^.

Existing methods to study membrane proteins in isolation from their cellular environments can be classified into supported bilayer (SB) and freestanding bilayer (FB) techniques. SB methods require forming a membrane on top of a solid (or gel) substrate^13–16^. Despite being easier to implement, these methods are inadequate for the study of IMPs because proteins are strongly affected by the presence of the substrate^17,18^. FB methods, where the membrane is suspended in solution, avoid immobilizing proteins, but current implementations of FBs have relied mostly on confocal optics that are not compatible with SPT^19–21^.

Here, we present the Freestanding-Bilayer Microscope (FBM) – a technique that combines the advantages of FBs with single-particle tracking. We based our instrument on planar lipid bilayers (PBs) for several reasons^22^. Most importantly, PBs have been shown to preserve the function of many ion channels of diverse types and organisms^23–26^. Also, PBs have diameters of hundreds of microns, which makes them ideal for imaging, and both sides of the membrane are accessible, allowing for great versatility in experimental design. Scanning a focused laser to illuminate the bilayer enabled us to observe fast diffusing IMPs at single-molecule resolution. We showed that in our system, IMPs diffuse freely as opposed to supported bilayers. We then showed how the FBM can be applied to answer diverse biological questions with two examples: the determination of the open probability of an ion channel by performing simultaneous imaging and electrical recordings, and the study of the dynamic organization of proteins and lipids by tracking diffusing IMPs in phase-separated lipid bilayers.

## Results

### Freestanding Bilayer microscope design

We designed the FBM to observe membrane proteins in PBs. PBs are formed by adding a small amount of lipid solution in an organic solvent, such as decane, across a hole in a plastic film (partition) submerged in an aqueous solution. The interfacial forces between the organic solvent, water, and the plastic material lead to the formation of a lipid bilayer at the center of the hole held in place by a Plateau-Gibbs border of lipid and solvent (torus) (Figure 1A). For the partitions, we used Fluorinated ethylene propylene (FEP) because it was the only material that fulfilled the conditions of low autofluorescence, good bilayer-forming properties, low refractive index, and transparency required for our experiments. Partitions were cut and perforated at the center, creating ∼200 μm diameter holes to support the bilayers, and were then mounted on a sample holder (cup) using vacuum grease to prevent leaks between the bottom and top chambers (Figure 1B). The cups attach to the experimental chamber, which is itself mounted on a micromanipulator and was designed to contain the tip of the excitation objective (XO) as well as ports for the perfusion of solution (bottom perfusion) (Figure 1B). The bottom perfusion ports are also connected to a manometer used to regulate the curvature of the bilayer (SI_Figure 1). Another set of perfusion channels was added to the cup to perfuse the top chamber (Figure 1B). To observe the bilayer under transmitted light, a glass window was included at the bottom of the chamber.

**Figure 1.**
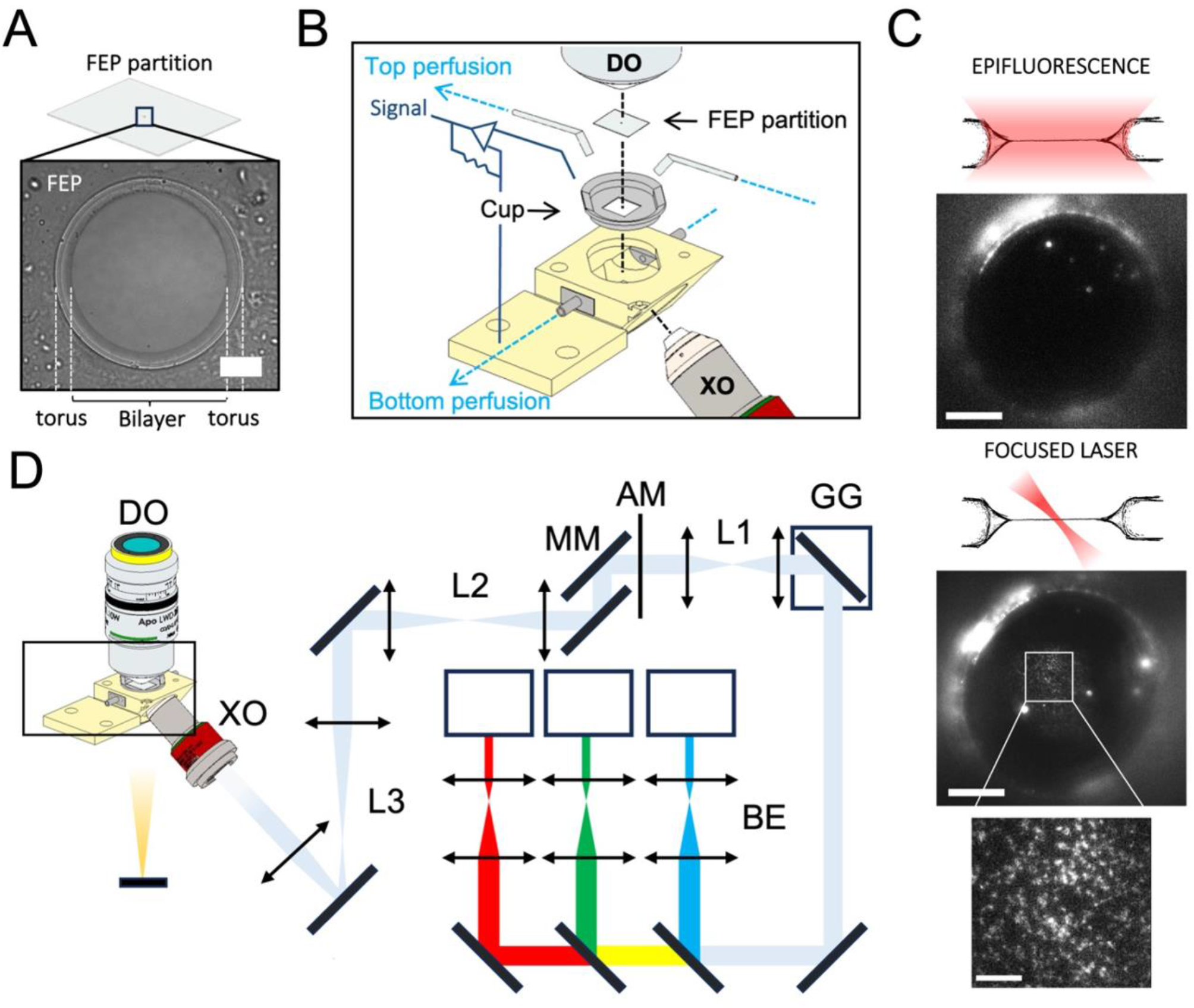
A. Image of a representative planar bilayer observed under transillumination. The boundaries between the bilayer, the Plateau-Gibbs border (torus), and the FEP partition can be seen. The bilayer composition is POPE: POPG (3:1 by weight). Scale bar: 50 μm. **B. Amplified-view of a technical illustration of the imaging chamber.** Planar bilayers are formed across perforated fluoroethylene- propylene copolymer films (FEP partitions) mounted on 3D-printed sample holders (Cup). Cups attach to the experimental chamber, shown in yellow, delimiting the top and bottom sides. The chamber provides access to the XO at 39 deg with respect to the optical plane of the bilayer, a transillumination source, and perfusion adaptors that can be connected to a perfusion system (bottom perfusion) or to a manometer to regulate the curvature of the bilayer. A different set of perfusion adaptors (top perfusion) are added to the top chamber along with the DO. Ag/AgCl electrodes are added to the top and bottom sides for electrophysiological recordings**. C. Fluorescent images taken with the FBM under epifluorescence (top) and focused-laser illumination (bottom) of fluorescently labeled hTRAAK.** Comparison between both illumination sources shows that resolved diffraction-limited particles can only be observed using focused-laser (FL) illumination. Insert shows a magnified view of the FL illumination. Scale bars: epi, 50 μm, FL, 50 μm, insert 10 μm. hTRAAK was labeled with LD655-NHS. **D. Scheme of the FBM depicting the relation between the main components of the system.** 473 nm, 561 nm and 660 nm lasers are fed to three beam expanders (BE) and optically conjugated to a galvo-galvo scanner (GG), an apodization mask (AM), and to the back focal plane of an excitation objective (XO) through three optical relays (L1-L3). Photons emitted from the sample are collected by a detection objective (DO).

Ag/AgCl electrodes were added to the top and bottom chambers to record currents across the bilayer to monitor the formation of the bilayer (sharp increase in capacitance) and to record the activity of ion channels in the bilayer (Figure 1B).

Since the PBs offer a natural optical sectioning of the emission of fluorescent proteins within it, we expected that an epifluorescence source would suffice to detect single molecules. However, the emission from the torus surrounding the bilayer caused a high background that prevented particle detection (Figure 1C, SI_Figure 2A, SI_Video 1). Inspired by the selective plane illumination microscopes (SPIM), we designed a scanned focused laser illumination that achieves precise illumination of fluorescent probes at the bilayer (Figure 1C and D, SI_Figure 2B)^27–29^. Focused illumination allowed us to observe single particles on the bilayer at typical frame rates of 50 Hz (Figure 1C, SI_Video 2). To achieve this, the lasers were first expanded and combined and then focused through a water immersion and long working distance objective at the bilayer (XO), positioned at an angle of 39 degrees with respect to the plane of the bilayer (Figure 1D and SI Figure 1). To define a small area of illumination within the bilayer, a galvo-galvo scanner (GG) and an apodization mask (AM) were conjugated to the back focal plane of the XO. By scanning the focused laser oblique to the bilayer at a higher frequency than acquisition, we created a virtual light sheet that produced a homogeneous illumination field (SI_Figure 2C). The AM contains concentric rings of different diameters that crop the incoming expanded laser to produce a Bessel beam after focusing it into the bilayer^28^. This creates a smaller and more stable focal point (SI_Figure 2B). Finally, a movable mirror (MM) was placed behind the AM to control the angle of illumination (Figure 1A) by changing the point of illumination on the back focal plane of the XO on the vertical axis. This increased the angle distribution by ±34 degrees.

**Figure 2.**
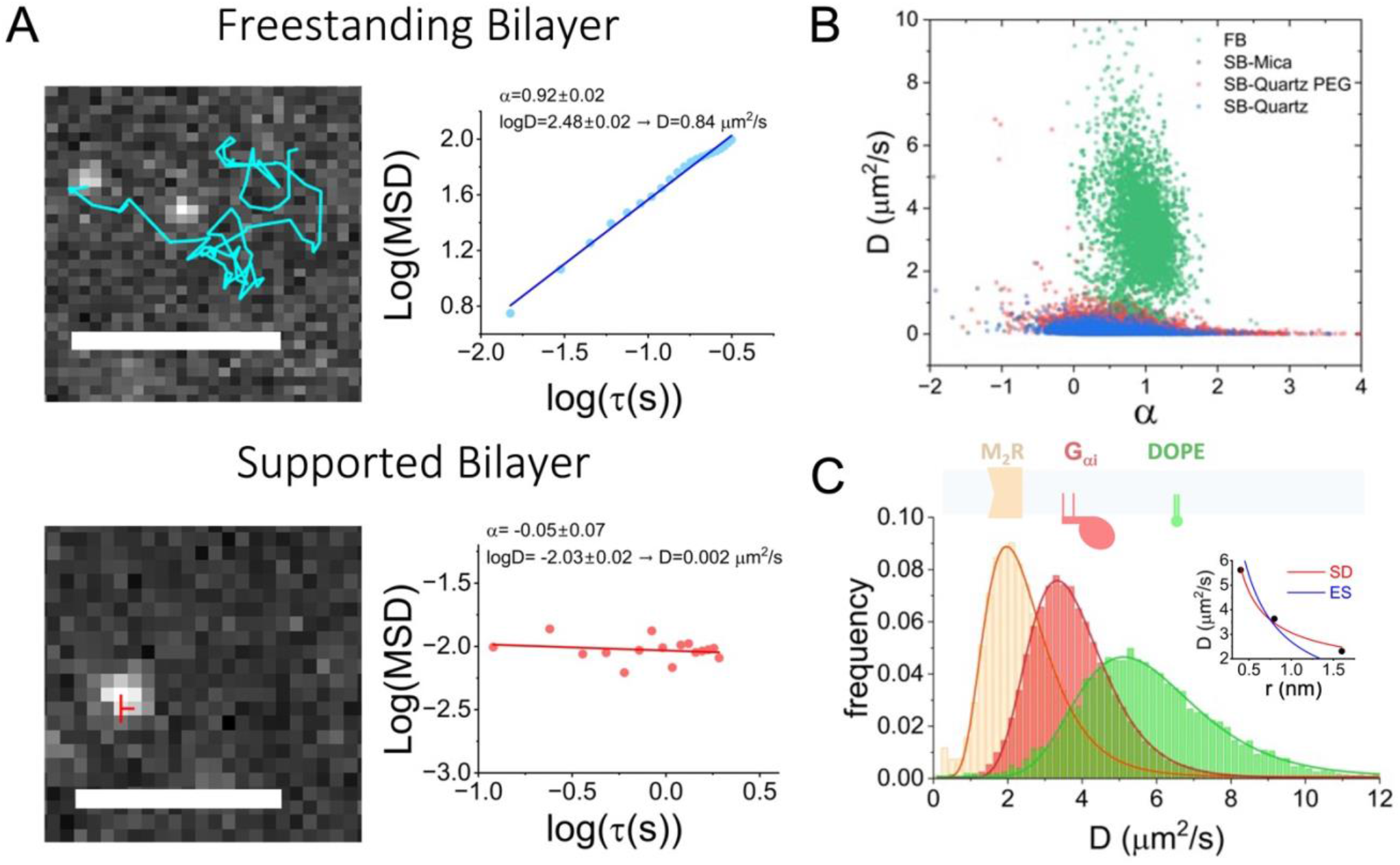
Single-Particle-Tracking benchmark of the FBM. A. Left: Representative SPT data for the M2R receptor diffusing in a freestanding bilayer (top) and in a Quartz supported bilayer (bottom). M2R receptor was reconstituted in POPE:POPG (3:1) vesicles for FBM experiments and in DOPC:POPG (7:3) for SBs experiments. For FBM experiments, vesicles were allowed to fuse into previously formed POPE: POPG (3:1) bilayers, followed by labeling with LD655-labeled NbALFA and extensive washing. For SBs, M2R-containing vesicles were allowed to burst and fuse onto solid substrates followed by washing. M2R was labeled with LD655-CoA through AcpS site-directed labeling. SB experiments were performed by TIRF illumination achieved with FBM optics, exchanging the FBM chamber for a prism- containing chamber. Scale bars: FBM: 5 μm, SB: 2 μm. **Right: MSD analysis of the tracks on the left.** Fit of the data indicates Brownian diffusion only for the FBM (top). **B. Diffusion (D) vs. anomalous diffusion (a) coefficients plots.** Data points were obtained from MSD analysis of 7620, 11307, 17731, and 6568 tracks for the FBM (green), mica-supported (grey), quartz-supported (blue), and pegylated- quartz-supported (red) membranes. The distribution of points for the FBM experiments (D ∼2 μm^2^/s and a=1±0.25) indicated that only FBM permits unconstrained diffusion of M2R. **C. Histrograms of D for M_2_R, G_αi1_ and DOPE-Cy5.** Experiments were performed using the FBM, observing unconstrained Brownian diffusion of all species. The distributions of D were fit to log-normal distributions (r^2^=0.99) and showed a weak dependence on size. G_αi_ was labeled with LD655 through Sfp site-directed labeling. Insert: Fit of the mean of distributions to a Saffman-Delbruck (r^2^ = 0.95) and Einstein-Smoluchowski (r^2^ = 0.68) model of diffusion.

The light emitted by fluorophores in the bilayer is collected by a commercial upright microscope coupled to a 4-camera splitter (SI_figure 1). To permit frequent and rapid access to the top chamber, we used water immersion objectives with long working distances. Because SPT requires both sufficient resolution and sufficient emitted light intensity, we found that, along with the focused illumination, the use of a 25X objective lens with high numerical aperture was critical to detect single molecules at fast frame rates.

### Diffusion of Integral Membrane proteins

To benchmark the performance of the FBM, we compared the diffusion of the human M2 Muscarinic receptor (M2R), an archetypical G-protein coupled receptor (GPCR) in our system against total internal reflection fluorescence (TIRF) experiments performed in supported bilayers (SB)^30^. SBs are relatively easy to implement in commercial microscopes, and thus they are extensively used as lipid membrane models^31–33^.

We formed SBs on three different substrates to account for any variability of the chosen substrate on the system: mica, quartz, and pegylated quartz^31,33^. Fluorescently labeled M2R was reconstituted in liposomes of DOPC: POPG (7:3)^34^. Liposomes were added to the substrate (coverslip) and incubated at 37 degrees to induce bursting and bilayer formation, followed by extensive washing before imaging by TIRF illumination. To perform TIRF experiments, we replaced the FBM experimental chamber with a prism-containing chamber (SI_Figure 3B). The coverslips were then mounted on the prism and imaged using the scanned focused laser of the FBM at total internal reflection (SI_Video 3-5). Videos were recorded at 8 Hz. For FBM experiments, we introduced an ALFA-tag at the C-terminus of M2R to render it amenable to effective and non-invasive fluorescent labeling after fusion to the planar bilayer^35^. This was done to avoid the increase in background produced by proteoliposomes partitioning into the torus. M2R was reconstituted in POPE:POPG (3:1) liposomes and fused into previously formed freestanding bilayers of the same lipid composition. After fusion, the top chamber was perfused with 5-10 chamber volumes to remove unfused vesicles, followed by the addition of 1 nM anti-ALFA nanobody (NbALFA) labeled with LD655. To remove excess NbALFA-LD655 the chamber was perfused with another 10 chamber volumes. The bilayers were then imaged using focused beam illumination, and videos were recorded at 50 Hz (SI_Video 6).

**Figure 3.**
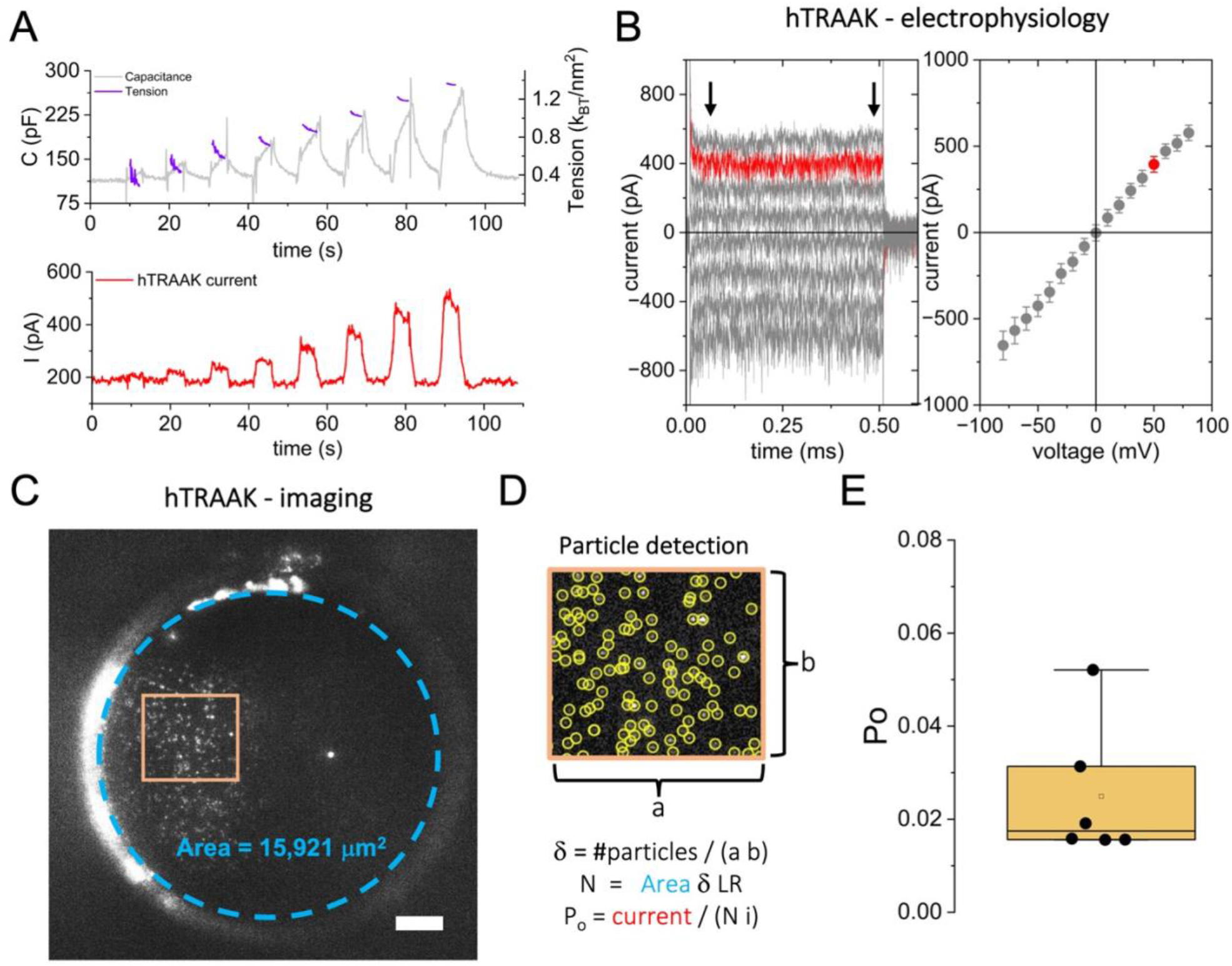
Simultaneous imaging and electrical recordings of ion channels. Determination of hTRAAK open probability (Po). A. Tension activation curves of hTRAAK showing its characteristic mechanosensitivity. The lateral tension of the bilayer (purple) was calculated from changes in the bilayers’ capacitance (grey) upon application of increasing pressure steps from 1 to 8 mmH2O (^40^). **B. Electrophysiology of hTRAAK.** Left: Potassium currents through hTRAAK channels, measured at voltage steps of 500 ms from -80 mV to 80 mV. Only odd voltage steps are shown for clarity. hTRAAK was reconstituted into POPE:POPG (3:1) vesicles and fused into bilayers of the same composition. Right: current-voltage characteristic showing the average ±SD of the currents at each voltage. The trace at 50 mV is plotted in red for illustrative purposes, and black arrows indicate the interval for the calculation of the average. **C. Imaging of hTRAAK.** frame of a video showing scanned focus-laser illumination of the hTRAAK containing bilayer from panel A. The dotted blue line shows the perimeter of the bilayer used to calculate its area. Scale bar: 20 μm. **D. Detection of hTRAAK tracks analyzed using SPT software.** Bottom: Equations used to calculate Po, where i is the single-channel current for hTRAAK and LR is the reciprocal of the labeled fraction of hTRAAK channels. **E. Po calculation for different bilayers.** The data are shown along a box plot showing the median (0.017), 25;75 percentile (box, 0.016;0.031), and 5;95 percentiles (whisker, 0.052) (N=6).

Videos for both SB and FBM experiments contained well-resolved diffraction-limited particles that were detected and analyzed using single-particle-tracking software^36^. Particle trajectories (tracks) were obtained after particle detection and assignment across successive frames. Figure 2A shows a characteristic example of a track for an FBM (top) and an SB (bottom) experiment. To analyze the tracking data, we plotted the mean squared displacement (MSD) of the track against increasing lag times (τ) and fit the data to the logarithmic anomalous diffusion equation,

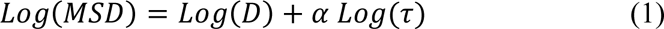

where D is the generalized diffusion coefficient and α is the anomalous coefficient, which takes a value of 1.0 ± 0.25 for normal (Brownian) diffusion^10,37^. We found that particles in SB are immobile, as evidenced by the shape of the MSD curve and the fitting results (D=0.002μm^2^/s, α=-0.05). Comparatively, particles in the FB are mobile, and the fit indicates Brownian diffusion (D=0.8μm^2^/s, α=0.92). We then performed a similar analysis on thousands of tracks (Figure 2B). The results, shown as D vs. α plots, reported a stark difference in the distribution of diffusion and anomalous coefficients; most particles undergo normal diffusion in the FBM whilst most particles are immobile in SBs regardless of the substrate material (Figure 2B and SI_Figure 4A). We reproduced these FBM experiments with two other membrane proteins with different folds and oligomeric structures, the dimer TRAAK and the tetramer GIRK2, finding similar results (SI_Figure 5), *i.e.* most particles (>80%) underwent Brownian diffusion with diffusivities ∼1.0 μm^2^/s.

**Figure 4.**
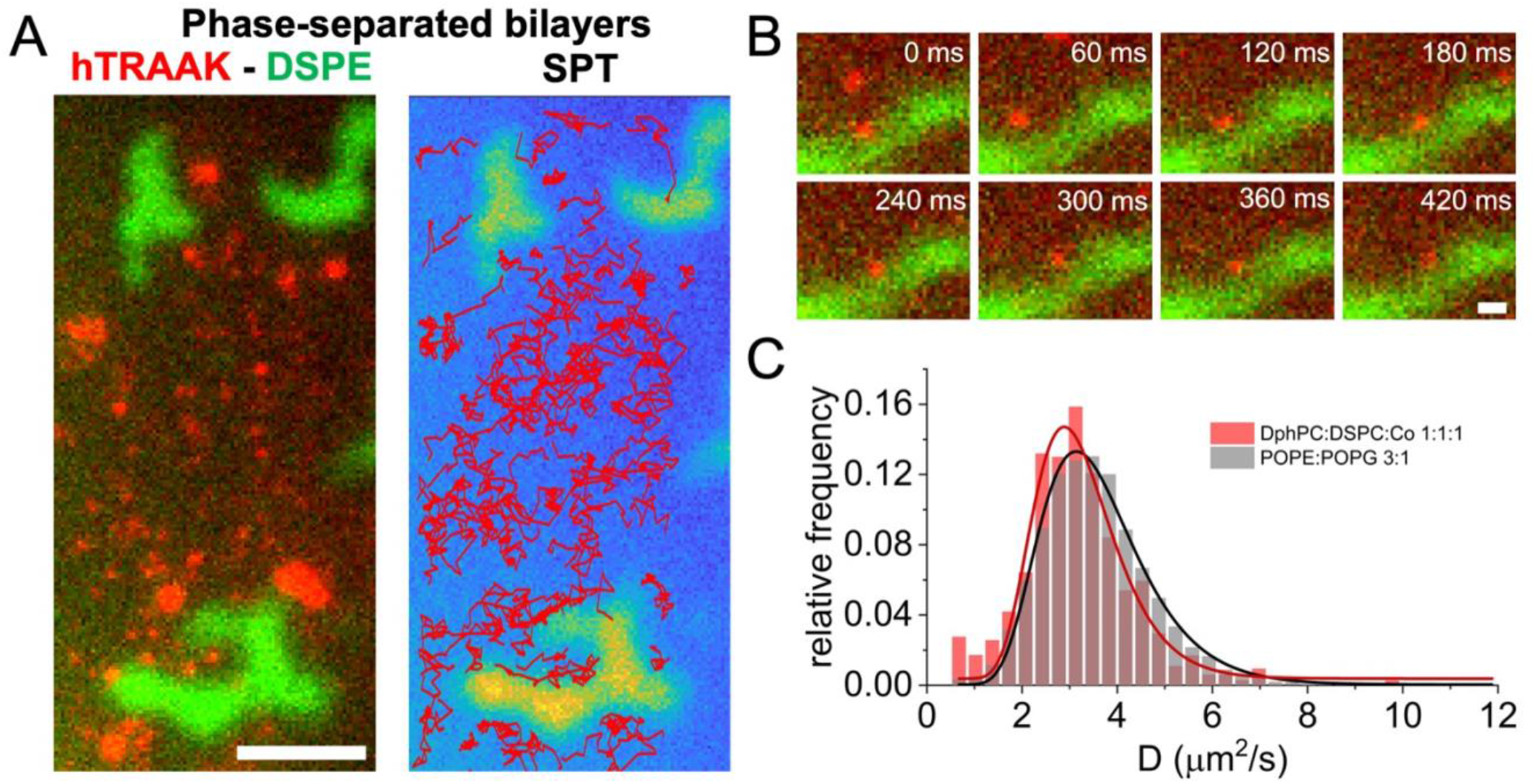
Single particle tracking of hTRAAK in phase-separated bilayers. A. Representative image of hTRAAK in DphPC: DSPC: cholesterol (2: 1 :1 molar ratio) Left: frame from a video overlapping hTRAAK (red) and DSPE-AF488 (green) channels. DSPE partitions into the ordered phase. Right: hTRAAK trajectories obtained from SPT analysis show the preference of hTRAAK for the disordered phase. Scale bar: 10 μm. hTRAAK was labeled with LD655-NHS. **B. Sequence of a single hTRAAK diffusing along the boundary between the two phases.** Scale bar: 2 μm **C. Diffusion coefficient (D) distributions of hTRAAK.** D were measured by SPT analysis in POPE:POPG (3:1 weight ratio, grey) and in DphPC: DSPC: cholesterol (2: 1 :1 molar ratio, red) bilayers showing no significant difference between the two lipid compositions.

Unconstrained Brownian diffusion was also observed for phospholipids (1,2-dioleoyl-sn-glycero- 3-phosphoethanolamine (DOPE-Cy5) and the fluorescently labeled membrane-associated guanine-nucleotide-binding protein G_αi_ (Supp. Videos 7-8). Analysis and classification of the tracks yielded histograms of diffusion coefficients that follow log-normal distributions (r^2^=0.99) (Figure 2C). We found that G_αi_ diffuses faster than M_2_R and the lipid DOPE faster than either protein. The relation between the diffusion coefficients of the three species is compatible with a model of diffusion that depends weakly on the hydrodynamic radius. Particularly, our data provide a better fit for a Saffman-Delbruck (SD) model (r^2^= 0.95) than for the Einstein-Smoluchowski (ES) model (r^2^=0.68) (Figure 2C)^38^.

Besides the substrates tested with SPT, we used FRAP to examine other solid support preparations (piranha-cleaned Quartz and SB formation on top of a Langmuir-Blodgett transferred lipid monolayer or over a transferred bilayer) to explore conditions that would possibly allow free diffusion^39^. In all cases tested, proteins in SBs were immobilized in contrast to FBs.

We should note that after the fusion of M2R-containing liposomes, we observed the presence of large aggregates of protein in the bilayer (SI_Figure 6A). These aggregates have somewhat round shapes and lie in the plane of the bilayer without noticeable height. We have observed similar aggregates for the ion channel GIRK2 (SI_Figure 6B) and TRAAK (Figure 4). Proteins exchanged between these aggregates and the rest of the bilayer. They also fused with each other in ways reminiscent of descriptions of protein condensates (SI_Figure 6C). We are currently studying these structures, and their description will not be part of the present publication.

### Applications of the FBM on ion channels. Estimation of open probability with multiple channel recordings

In FB experiments, both sides of the bilayer are isolated, which permits coupling single-particle imaging experiments with functional measurements that require transport across the bilayer. We used this property of the FBM to determine the open probability of an ion channel in bilayers containing multiple channels using simultaneous imaging and electrical recordings. To perform these experiments, we chose the human mechanosensitive potassium channel TRAAK (hTRAAK). We have previously studied hTRAAK in freestanding bilayers, showing tension sensitivity curves measured with a freestanding bilayer tensiometer that we developed and which is fully compatible with the FBM^40^ (Figure 3A).

Determining ion channel open probability (Po) can usually be accomplished by measuring single channels or performing noise analysis on recordings with multiple channels, so called macroscopic current recordings^41^. However, the single-channel kinetics of TRAAK and its very low open probability, which renders a linear relationship between current variance and mean, preclude a good estimation of Po by either of the aforementioned methods.

The exact number of channels that contribute to the current is the main unknown in the effort to determine the Po of a channel from macroscopic experiments. Using the FBM, we were able to determine the number of hTRAAK channels (N) in the bilayer while simultaneously measuring the current through those channels (I) (Figure 3B, C and D). Knowing N and I, we calculated hTRAAK Po from the simple relation, I = i N Po, where i is the current through a single channel^42^. We determined the median of the channel Po at the bilayer basal tension (0.2-0.3 kBT/nm^2^) to be 1.7% (IQR=1.6;3.1) (Figure 3E). Furthermore, increasing the tension of the bilayer by about 4-fold increased the channel activity by only 2.5-fold.

For these experiments we purified, labeled, and reconstituted hTRAAK in POPE:POPG (3:1, weight ratio) liposomes. After liposome fusion into FBs of the same lipid composition, we recorded videos of diffusing channels and performed SPT analysis as described above. From this analysis, we calculated the density of detected particles within a region of the bilayer with homogeneous focus and illumination and extrapolated the number of hTRAAK in the entire bilayer after correcting for the labeling efficiency and subsequent multiplying by the total bilayer area (Figure 3D).

### Protein partitioning into phase-separated bilayers

Cell membranes contain hundreds of different kinds of lipids^43^. It has been shown that the interaction between different species of lipids can give rise to critical phenomena. For example, ternary mixtures of lipids of a lipid with a high Tm, a lipid with low Tm, and cholesterol can phase-separate in vitro, leading to the coexistence of liquid-ordered and liquid-disordered phases^44^. Additionally, it has been observed that the protein distribution in membranes is not homogeneous, and some have proposed that inhomogeneous lipid composition could be a driving mechanism of this distribution^45^. To test whether lipid composition and phase-separation can drive an inhomogeneous distribution of proteins, we performed single-particle-tracking experiments with hTRAAK in bilayers of DphPC:DSPC:cholesterol (2:1:1), a mixture that phase separates at room temperature, forming liquid ordered and liquid disordered phases^46^. We added a small amount of fluorescent lipid to label the ordered phase (DSPE-AF488). The data show a striking preference of hTRAAK for the disordered phase (Figure 4A). This preference is further evidenced by the distribution of tracks that avoid the ordered regions (Figure 4A, right panel). TRAAK molecules that approached the boundary between phases often moved along the perimeter before returning to the disordered phase (Figure 4B). We also found that the diffusion coefficient distribution of TRAAK in the phase-separated bilayer does not differ significantly from the distribution in the binary mixture POPE:POPG (3:1) (Figure 4C).

## Discussion

Integral membrane proteins are fundamental elements of cell signaling and energy metabolism. Yet, they remain incredibly difficult to study due to their low expression levels and the complexity of their native environments. Current techniques cannot fully address the dynamics of proteins in membranes; how they interact with themselves and others, and how their motion and distribution are affected by the environment.

We introduced FBM, Freestanding Bilayer Microscope, a method to perform in vitro single-molecule imaging of membrane proteins in membranes. We showed that the use of planar freestanding bilayers, as opposed to supported bilayers, leads to unconstrained free diffusion devoid of artifacts caused by the presence of the substrate. We demonstrated the use of the FBM by performing single-particle-tracking of the integral membrane protein M_2_R, as well as the lipidated G_αi1_ and fluorescent DOPE lipids. From the tracking results, we found that over 90% of the tracks underwent Brownian diffusion. We determined the distribution of diffusion coefficients for the three species, finding that they fit to a log-normal distribution. By fitting the median of the distributions to the estimated hydrodynamic radius of each molecule, we found that the data correspond better to a Safmann-Delbruck model than to an Einstein-Smoluchowski model, which implies a dependence of diffusion on radius weaker than (radius)^-138^.

The FBM is based on planar bilayers (PB), a proven, effective method for the fully functional reconstitution of different membrane proteins utilized in different laboratories worldwide. This offers versatility as the FBM setup can easily be combined with other PB-based assays. We demonstrated this by performing simultaneous imaging and electrical recordings of the mechanosensitive channel hTRAAK. hTRAAK is located in the Nodes of Ranvier of myelinated neurons of mammals, and it has been shown to play an important role in thermal and mechanical nociception^47^. Determining the open probability, Po, of this channel has been challenging owing to its flickery single-channel kinetic behavior. By using the FBM, we determined that the Po of hTRAAK at the basal tension of PBs is 2.5%, with an estimated increase to 6.3% at tensions up to 1.2 kBT/nm^2^, which is about 60% of the lytic tension needed to rupture a typical lipid membrane. These Po values imply that hTRAAK works within a range of low Po and are consistent with previous results from our lab^48^.

We also showed that the FBM can be used to study the influence of lipid composition on the diffusion and distribution of membrane proteins. We explored this possibility using a ternary mixture of lipids that phase separate at room temperature. Incorporation of TRAAK into these bilayers did not change the amount of lipid phase separation, but there was a striking preference of TRAAK for the liquid disordered (Ld) phase. No particles of TRAAK were detected within the ordered phase. This phenomenon suggests a mechanism of exclusion and enrichment of TRAAK that could play an important role in the much more complex plasma membrane of cells.

We anticipate that the FB microscope will be an important tool to elucidate the mechanisms of membrane protein organization and dynamics. For example, the information contained in single-particle tracking can be used to study protein-protein interactions in the membrane. Combined with the unique compositional control offered by the planar bilayers, this could lead to new insights into signaling mechanisms that depend on dynamic complex formation, such as G-protein coupled receptors, receptor tyrosine kinases, or immune receptor pathways. Furthermore, the possibility of combining the FBM with smFRET offers the possibility to study structural changes of proteins as a function of interactions with other species in a reaction-diffusion setting, something that is today only possible with simulations.

Finally, our results show that the diffusion of membrane proteins in FBs is different than in cell membranes. Diffusion coefficients in FBs are larger than in cell membranes by about ten-fold, and a larger fraction of the diffusion is Brownian. There are many hypotheses to explain smaller diffusion coefficients, higher immobile fractions, and anomalous diffusion in cell membranes^4,49,50^. The FBM provides a good starting point to test these hypotheses experimentally. The FBM is a reductionist system that will permit controlled, stepwise building in of complexity so that we may eventually understand the properties of diffusion in cell membranes, and ultimately the signaling processes that rely on cell membrane diffusion.

## Materials and methods

### M2R-ALFA and A1-M2R expression and purification

An ALFA-tag was introduced at the C terminus of the human muscarinic 2 receptor (hM2R) gene, followed by a 3C PreScission Protease (PPX) recognition site, the eYFP gene, and a polyhistidine tag (His10-tag) (M2R-ALFA). The ALFA-tag was used for binding with an anti-ALFA nanobody (ALFANb) conjugated to a fluorophore for single molecule visualization, while the eYFP and His10-tags were inserted for purification purposes^35^. The gene was then cloned into a pFastBac vector for protein expression with the Bac-to-Bac Baculovirus System. *Spodoptera frugiperda* Sf9 insect cells were transfected with isolated bacmid DNA to produce a recombinant baculovirus stock. The virus was amplified through two more rounds of progressive infection of Sf9 cells to create a high-titer stock used for large-scale expression of M2R-ALFA. Six liters of Sf9 cells were cultured to a density of 3 million cells per mL and infected with 20 mL of baculovirus stock. The cells were grown for 48 hours post-infection at 27℃ shaking at 120 rpm, after which cells were pelleted by centrifugation at 3,500g for 15 minutes at 4℃. Cell pellets were resuspended in 1x phosphate-buffered saline (1x PBS; Thermo Fisher Scientific), and centrifuged at 3,500g for ten minutes at 4℃. Pellets were flash-frozen in liquid nitrogen and stored at -80℃ until use.

A short peptide tag, A1, was cloned at the N terminus of the hM2R gene to enable post-purification AcpS phosphopantetheinyl transferase (PPTase) catalyzed site-specific protein labeling (A1-M2R) (^51^). At the C terminus of the hM2R gene, the PPX recognition site, eYFP gene, and His10-tag were included for purification purposes. A1-M2R was expressed as described for M2R-ALFA.

M2R constructs (M2R-ALFA and A1-M2R) were purified following the same protocol^52^. All purification steps were carried out at 4℃ unless specified otherwise. Sf9 cells were lysed by osmotic shock, by resuspending the pellets in the following buffer for 30 minutes: 10 mM Tris HCl pH 7.4, 10 mM MgCl2, 5 units per mL benzonase nuclease, 5 mM β-mercaptoethanol (βME), 2 mM phenylmethylsulfonyl fluoride (PMSF), 1x of 3-7 TIU/mg aprotinin saline solution, and 0.1% v/v of protease inhibitor cocktail (PIC) stock (1 mg/mL leupeptin, 1 mg/ml pepstatin A, 1 M benzamidine HCl). After mixing for 30 minutes, the lysis slurry was centrifuged at 30,000g for 20 minutes. Once the supernatant was removed, cell pellets were resuspended in high salt buffer (20 mM HEPES pH 7.4, 750 mM NaCl, 10% v/v glycerol, 5 mM βME, 5 units per mL benzonase nuclease, 5 µM atropine, 1x of 3-7 TIU/mg aprotinin saline solution, 0.1% v/v of PIC stock) and homogenized using a dounce homogenizer. IMPs were extracted by solubilizing the membrane with 1%:0.05% n-dodecyl-β-D-maltopyranoside: cholesterol hemisuccinate (DDM:CHS) for one hour at room temperature, followed by one hour at 4℃.

After extraction, insoluble material was pelleted by centrifugation at 30,000g for 30 minutes. The supernatant was bound to TALON resin and washed with buffers (20 mM HEPES pH 7.4, 0.1%:0.01% DDM:CHS, 1 µM atropine, 1mM βME, 0.1% v/v of PIC stock) containing a gradual increase in NaCl concentration. The resin was washed with five column volumes (CV), referring to the volume of resin used, of wash buffer containing 750 mM and 587 mM NaCl, respectively. Next, the resin was washed with five CVs of wash buffer with 424 mM NaCl and 20 mM imidazole. M2R was eluted with 261 mM NaCl and 200 mM imidazole wash buffer.

The elution fractions were pooled and subsequently diluted tenfold with the following dilution buffer in order to lower the imidazole concentration and introduce the superagonist iperoxo: 20 mM HEPES pH 7.4, 100 mM NaCl, 0.1%:0.01% DDM:CHS, 0.1 mM TCEP, 12 µM iperoxo. The protein was then bound to anti_GFP nanobody-coupled Sepharose resin (GFPNb resin), produced in-house as previously described, for 1 hour^52^. The protein was buffer exchanged by washing the resin on the gravity column with 20 CV of buffer containing the same components as the dilution buffer, except for an increased iperoxo concentration of 50 µM. PPX was added to the resin. The protein was nutated overnight to ensure complete digestion and to change the ligand bound to M2R from the antagonist atropine to iperoxo. The cleaved M2R was eluted by collecting the flow through of the GFPNb resin by gravity.

A1-M2R was labeled with two-fold excess LD655-CoA dye in the presence of 10 mM MgCl2, 2 mM TCEP, and a 1:5 ratio of AcpS (expressed and purified in-house) to A1-M2R. The enzyme catalyzed fluorophore conjugation reaction was allowed to proceed in the dark for 2 hours at room temperature, followed by 2 hours at 4℃. A labeling efficiency of ∼10% was attained using this method. M2R-ALFA was fluorescently labeled through binding with NbALFA-LD655 after vesicle fusion into freestanding bilayers.

M2R constructs were dephosphorylated through a 30-minute room temperature treatment with 200 units of lambda protein phosphatase (NEB), 10 units of calf intestinal phosphatase (NEB), 5 units of antarctic phosphatase (NEB), and 1 mM manganese chloride. Proteins were concentrated in 30 kDa MWCO concentrators prior to their respective purification by size exclusion chromatography using a Superdex 200 Increase 10/300 GL column and running buffer (20 mM HEPES pH 7.4, 100 mM KCl, 50 mM NaCl, 10 µM iperoxo, 100 µM TCEP, 0.1%:0.01% DDM:CHS). M2R reconstitution in lipid vesicles was done immediately after gel filtration. Purified M2R that was not reconstituted was flash frozen with 10% glycerol and stored at -80℃.

### G_αi1_ expression and purification

The short peptide tag, S6, was inserted in an internal loop between amino acids 112 and 113 of the human Gɑi1 gene^51^. The S6 tag was inserted for site-specific protein labeling via the Sfp PPTase enzyme catalyzed peptide tag modification. The Gɑi1-S6 gene was cloned into the pFastBac vector. Genes for the wild-type human Gβ1 and Gγ2 subunits were also cloned into the pFastBac vector. At the N terminus of Gγ2, a His10-eYFP linked to a PPX recognition site was inserted. Recombinant baculovirus for each subunit was prepared separately as described above. For large-scale expression of the heterotrimeric G protein, six liters of *Trichoplusia ni* High Five insect cells were cultured to a density of 2 million cells per mL. Each liter of cells was coinfected with an experimentally determined ratio of Gɑi1, Gβ1, Gγ2 baculovirus (8 mL of Gγ2, 12 mL of Gβ1, and 15 mL of Gɑi1) to achieve expression of the heterotrimeric G protein subunits in stoichiometric proportions. The cells were grown for 36-48 hours and then harvested by centrifugation at 3,500g for 15 minutes at 4℃. Cell pellets were resuspended in 1x PBS and centrifuged at 3,500g for 10 minutes. Pellets were flash-frozen and stored at -80℃ until further use.

All purification steps were carried out at 4℃ unless specified, and stock guanosine 5’-diphosphate disodium salt (GDP) aliquots were made fresh before use. Cell pellets were resuspended in buffer containing 20 mM Tris HCl pH 8, 100 mM NaCl, 5 mM DTT, 3 mM MgCl2, 20 µM GDP, 2 mM PMSF, DNase I, 1x of 3-7 TIU/mg aprotinin saline solution, and 0.1% v/v of PIC stock. After mixing for 15 minutes, the cell slurry was manually homogenized. Membranes were extracted in 1% sodium cholate for 1.5 hours. The extraction was centrifuged at 35,000g for one hour. The supernatant was bound to GFPNb resin and then washed with 10 CVs of wash buffer (20 mM Tris HCl pH 8, 100 mM NaCl, 1% Na-cholate, 3 mM MgCl2, 20 µM GDP) on column. Gɑi elution buffer (20 mM Tris HCl pH 8, 100 mM NaCl, 1 % Na-cholate, 50 mM MgCl2, 30 µM GDP, 10 mM NaF, 30 µM AlCl3) was used to elute Gɑi1-S6 from the G_βγ_ complex bound to the GFPNb resin. The resin was moved to room temperature, and the elution buffer was warmed in a 30℃ water bath for 15 minutes; GDP was added to the elution buffer afterward. The resin and elution buffer were incubated together, nutating for 30 minutes at room temperature, and Gɑi1- S6 alone was collected by gravity flow. The eluate containing Gɑi1-S6 was labeled using two-fold molar excess LD655-CoA dye, Sfp enzyme that was purified in-house (1:2 ratio of enzyme to protein), and 10 mM MgCl2. A labeling efficiency of ∼10% was obtained. Gɑi1 was dephosphorylated through a 30 minute room temperature treatment with 200 units of LPP, 10 units of CIP, 5 units of AP, and 1 mM manganese chloride. The protein was concentrated and purified using a Superdex 200 Increase 10/300 GL column (20 mM Tris HCl pH 7, 150 mM KCl, 10 mM DTT, 0.4 mM TCEP, 1% Na-cholate, 2 mM MgCl2, 20 µM GDP). Fractions containing LD655-Gɑi were collected and reconstituted into lipid vesicles. Non-reconstituted protein was flash-frozen with 10% glycerol and stored at -80℃.

### hTRAAK expression and purification

The human mechanosensitive potassium channel TRAAK (hTRAAK) gene was cloned into a modified pPICZ-B vector to create a fusion protein construct containing a C terminal PPX cleavage site, the GFP gene, and His10-tag. The hTRAAK protein construct was then transformed into *P. pastoris* strain SMD11-63H and expressed in large scale cultures as described previously^53^. Cell pellets were flash frozen in liquid nitrogen and subsequently disrupted by milling 5 times for 3 minutes at 25 Hz. Lysed cells were frozen and stored at -80℃ until they were ready for purification.

All purification steps were carried out at 4℃. The lysed cells were mixed with a five-fold excess volume of buffer (50 mM Tris HCl pH 8.5, 150 mM KCl, 1 mM EDTA, DNase I, 1x of 3-7 TIU/mg aprotinin saline solution, 0.1% v/v of PIC stock) relative to the wet pellet volume for 15 minutes. The membranes were manually homogenized and extracted using 2% DDM for 1.5 hours. The extraction mixture was centrifuged at 39,000g for 30 minutes. The supernatant was bound to GFPNb resin for one hour. The resin was washed with 10 CV of wash buffer (20 mM HEPES pH 8, 150 mM KCl, 0.1% DDM). PPX was added to the resin in a 1:20 protease to protein ratio, and the protein was digested overnight. The flow through containing cleaved TRAAK was collected from the resin and was concentrated using a 10 kDa MWCO concentrator.

TRAAK was non-specifically labeled using amine-reactive LD655-NHS dye. The protein was buffer exchanged using a PD-10 column into 1x PBS without calcium and magnesium at pH 8. 50-fold molar excess LD655-NHS dye was added to the protein solution; protected from light, the reaction was allowed to proceed overnight at 4℃. The protein was labeled with an efficiency of 15%. Labeled TRAAK was concentrated using a 10 kDa MWCO concentrator and subsequently purified using a Superdex 200 Increase 10/300 GL column (20 mM Tris-HCl pH 7.4, 150 mM KCl, 1 mM EDTA, 0.03% DDM). Fractions containing LD655-TRAAK were reconstituted.

### GIRK-ALFA expression and purification

At the C terminus of hGIRK2, a PPX cleavage site, eGFP gene, and His10 tag were preceded by an ALFA-tag. The fusion protein construct was cloned into the pBacMam vector and the BacMam system was used to create a high titer recombinant baculovirus stock using Sf9 cells. Four liters of the HEK293S GnTI^-^ strain from ATCC (CRL-3022) were cultured to a density of 3.5 million cells per mL and infected with 100 mL of the high titer GIRK2-ALFA recombinant baculovirus. After 12 hours, the cells were induced with 10 mM sodium butyrate. After shaking at 37℃ for an additional 40-48 hours, the cells were harvested by centrifugation at 3,500g for 15 minutes at 4℃. Cell pellets were resuspended in 1x PBS and centrifuged at 3,500g for ten minutes at 4℃. The supernatant was discarded. Cell pellets were flash frozen and stored at -80℃.

Purification of GIRK2-ALFA was carried out at 4°C unless noted otherwise. Cell pellets were first mixed in resuspension buffer (25 mM Tris HCl pH 7.5, 150 mM KCl, 50 mM NaCl, 1 mM MgCl2, 1 mM CaCl2, 2 mM DTT, 2 mM PMSF, DNase I, 1x of 3-7 TIU/mg aprotinin saline solution, 0.1% v/v of PIC stock) for 15 minutes. The cell slurry was homogenized (Dounce), and the lysate was centrifuged at 39,000 xg for 15 minutes. The supernatant was discarded. Pellets were suspended in buffer (25 mM Tris HCl pH 7.5, 150 mM KCl, 50 mM NaCl, 1 mM MgCl2, 1 mM CaCl2, 2 mM DTT, 1x of 3-7 TIU/mg aprotinin saline solution, 0.1% v/v of PIC stock) and homogenized (Dounce). Membranes were extracted with 1.5%:0.3% DDM:CHS for 2 hours. The extraction was centrifuged at 39,000g for 30 minutes to pellet the insoluble fraction, and the supernatant was bound to GFPNb resin for one hour. The resin was washed on column with 20 CVs of wash buffer (20 mM Tris HCl pH 7.5, 150 mM KCl, 50 mM NaCl, 2 mM DTT, 0.05%:0.01% DDM:CHS). PPX was added to the resin slurry for overnight digestion. The flowthrough containing cleaved GIRK2-ALFA was collected from the GFPNb resin by gravity flow. The protein was concentrated using a 100 kDa MWCO concentrator and purified of contaminants using a Superose 6 Increase 10/300 GL column with the buffer: 20 mM Tris HCl pH 7.5, 150 mM KCl, 50 mM NaCl, 10 mM DTT, 1mM Na2EDTA pH 7.4, 0.025%:0.005% DDM:CHS. GIRK2-ALFA was reconstituted into liposomes after gel filtration. The GIRK2-ALFA construct was fluorescently labeled with NbALFA- LD655 after vesicle fusion into freestanding bilayers.

### Anti-ALFA Nanobody expression and purification

The NbALFA was expressed as an N terminal His14-SUMO fusion in *E. coli* cells^35^. The recombinant plasmid, was transformed into One Shot BL21 Star (DE3) cells. A single colony was used to inoculate small-scale cultures: 50 mL of lysogeny broth (LB) with 50 µg/mL of kanamycin. The cells were grown overnight (∼ 18 hours) at 37℃ shaking at 225 rpm. The next morning, 20 mL of small-scale culture was added to one liter of LB supplemented with 50 µg/mL of kanamycin for a 1:50 dilution. Once the cells reached an optical density (OD) at 600 nm of about 0.6, the cells were induced with 0.5 mM IPTG and allowed to grow overnight at 16℃. Cells were harvested by centrifuging at 3,500 g for 15 minutes at 4℃. Cell pellets were resuspended in 1x PBS, and centrifuged at 3,500 g for 10 minutes at 4℃. Flash frozen pellets were stored at -80℃.

All purification steps were carried out at 4°C unless otherwise specified. A cell pellet was suspended in lysis buffer (20 mM HEPES pH 7.9, 300 mM NaCl, 2 mM PMSF, DNase I, and 1 mM TCEP, 2x of 3-7 TIU/mg aprotinin saline solution, 0.2% v/v of a protease inhibitor mixture stock [0.1 g/mL trypsin inhibitor, 1 mg/mL pepstatin A, 1 mg/mL leupeptin, 1 M benzamidine HCl, 0.5 M AEBSF] and mixed for 15 minutes. The cells were lysed by sonication and the lysate was clarified by centrifuging at 16,500 rpm for 40 minutes at 4℃. The supernatant was bound to equilibrated Ni-NTA resin for one hour and washed with wash buffer (20 mM HEPES pH 7.9, 300 mM NaCl, and 1 mM TCEP) containing no imidazole followed by 20 mM imidazole. ALFANb was eluted off the resin with 400 mM imidazole containing wash buffer. The His14-SUMO tag was cleaved of the protein by adding ULP1 (prepared in-house) to the elution. The NbALFA-protease solution was placed in 8 kDa MWCO dialysis tubing and dialyzed overnight at room temperature against a buffer composed of 20 mM HEPES pH 7.9, 300 mM NaCl, 0.5 mM TCEP, and 2 mM DTT. The insoluble precipitate was removed by centrifuging the protein for 10 minutes at 3,500g. The supernatant was run through Ni-NTA resin equilibrated in wash buffer containing 10 mM imidazole in order to collect cleaved NbALFA. The digested protein was concentrated with a 10 kDa MWCO concentrator and purified using a Superdex 75 10/300 GL column (20 mM HEPES pH 7.9, 150 mM NaCl, 1 mM TCEP). Purified NbALFA was stored at -80°C.

To prepare fluorescent NbALFA for binding ALFA tagged IMPs on the bilayer, the cysteines of purified NbALFA were non-specifically labeled with maleimide LD655 (LD655-MAL). Thawed NbALFA was incubated with 0.1 mM of freshly made TCEP at room temperature for 10 minutes. The reduced protein was buffer exchanged into 1x PBS (not containing calcium and magnesium) at pH 7-7.5 using a PD-10 desalting column to remove excess TCEP, which has been found to interfere with high efficiency bioconjugation by reacting with maleimides to form other products. The NbALFA was mixed with five-fold molar excess LD655-MAL and reacted overnight in the dark at 4°C. To separate the labeled nanobody from excess dye, the protein was run on the Superdex 75 10/300 GL column equilibrated with 20 mM HEPES pH 7 and 150 mM NaCl. A labeling efficiency of ∼80% was achieved; NbALFA-LD655 was stored at -80°C.

### Proteoliposome (PLs) reconstitution

Membrane proteins were reconstituted into liposomes immediately after labeling (if applicable) and purification. A given protein was reconstituted using a specific mixture of phospholipids. The following general protocol to dry lipid films was repeated for all lipid combinations used to reconstitute the proteins discussed in this paper. The desired ratio of lipids in chloroform was mixed in a glass vial, and the chloroform was evaporated under a steady stream of argon gas. The lipid film was solubilized in a small amount of pentane and the solvent was again evaporated under a steady stream of argon. The lipid film was thoroughly dried by storage in a vacuum desiccator overnight.

For FBM experiments, M2R-ALFA was reconstituted into a 3:1 weight ratio of 1-palmitoyl-2- oleoyl-sn-glycero-3-phosphoethanolamine (POPE): 1-palmitoyl-2-oleoyl-sn-glycero-3-phospho-(1’-rac-glycerol) (POPG) lipids. After removal from the desiccator, the lipid film was rehydrated in buffer (20 mM HEPES pH 7.4, 100 mM KCl, 50 mM NaCl, 10 µM iperoxo, 100 µM TCEP) to a stock concentration of 10 mg/mL. The POPE:POPG (3:1) lipids were sonicated to clarity in a water bath sonicator. 1% n-Decyl-β-maltoside (DM) was added to the lipids to foster protein insertion into vesicles. The lipids were nutated for 30 minutes at room temperature and sonicated again. To create a protein to lipid ratio (PLR) of 1:20, M2R-ALFA and POPE:POPG lipids were combined to a final working concentration of 0.25 mg/mL and 5 mg/mL, respectively. The mixture was nutated for one hour at room temperature. To remove the detergent, biobeads (Bio-Beads SM-2 Adsorbent, Bio-Rad) equilibrated in 20 mM HEPES pH 7.4, 100 mM KCl, 50 mM NaCl, 10 µM iperoxo, and 100 µM TCEP were added to the protein-lipid mixture so that the dry volume of beads was roughly one-third the volume of vesicles. The mixture was moved to nutate at 4°C and biobeads were changed every 8-12 hours for a total of 3-4 changes. After detergent removal, the proteoliposomes were harvested, flash frozen and stored at -80°C.

A1-M2R was reconstituted into a 7:3 weight ratio of 1,2-dioleoyl-sn-glycero-3-phosphocholine (DOPC): POPG lipids for SB experiments. The same reconstitution process and buffers were used as described above for M2R-ALFA. The protein was protected from light during reconstitution to prevent bleaching of the LD655 fluorophore conjugated to the receptor.

Gɑi1-S6 was reconstituted into a 3:1 weight ratio of POPE:POPG lipids. The lipids were brought to a concentration of 10 mg/mL using 20 mM Tris HCl pH 8, 150 mM KCl, 2 mM MgCl2 and sonicated to clarity in a water bath sonicator, followed by addition of 1% DM and room temperature incubation for 30 minutes. After more sonication, a mixture with 0.405 mg/mL Gɑi1-S6 and 5 mg/mL lipids was made for a PLR of 1:12. The mixture was nutated for one hour at room temperature in the dark. Dialysis was then used to remove the detergent. The mixture was placed in 10 kDa MWCO dialysis tubing, and the buffer (20 mM Tris HCl pH 8, 150 mM KCl, 20 mM MgCl2, and 5 mM DTT) was exchanged five times every 12 hours, adding fresh DTT each time. Proteoliposomes were harvested, flash frozen and stored at -80°C.

For FBM experiments, GIRK2-ALFA was reconstituted into a 3:1 weight ratio of POPE:POPG lipids. Lipids dried overnight were made 20 mg/mL using buffer (20 mM Tris HCl pH 7.5, 150 mM KCl, and 50 mM NaCl). Lipids were sonicated to clarity, and 1% DM was added. After lipids were incubated at room temperature for 30 minutes, they were sonicated again and combined with 2 mg/mL GIRK2- ALFA in equal parts to create a final PLR of 1:10. Following a one-hour nutation at room temperature, the vesicles were placed in dialysis tubing of 50 kDa MWCO and dialyzed in 10 mM K2HPO4 pH 7.4, 150 mM KCl, and 1 mM K2EDTA. The buffer was exchanged five times every 12 hours, adding fresh DTT for each change. For the last two buffer exchanges, equilibrated biobeads were included in the buffer. Proteoliposomes were aliquoted, flash frozen and stored at -80°C.

TRAAK was also reconstituted into a 3:1 weight ratio of POPE:POPG. Lipids were dissolved at 10 mg/mL in 20 mM Tris HCl pH 7.4 and 150 mM KCl buffer. The lipids were sonicated to clarity. 1% DM was added, and the lipid mixture was nutated at room temperature for 30 minutes, followed by further sonication. A PLR of 1:10 was achieved by combining channel and lipids for a final concentration of 0.5 mg/mL and 5 mg/mL, respectively. After one hour of incubation at room temperature, the vesicles were dialyzed in 30 kDa MWCO tubing using 10 mM Tris HCl pH 7.4 and 150 mM KCl buffer containing equilibrated biobeads. The buffer was exchanged every 12 hours for a total of three times. The vesicles were then removed from dialysis tubing and mixed directly with biobeads for 6 hours prior to being harvested, flash frozen, and kept at -80°C until use.

### Freestanding bilayer Microscope (FBM) set-up. Detection optics

Data acquisition was done with up to 4 sCMOS cameras (Orca-Flash4.0 V3, Hamamatsu) connected to a commercial upright microscope (FN1, Nikon Instruments), with four 10 mm spacers between the arm and the head, (FN-S10, Nikon Instruments) through a camera splitter (Multicam LS image splitter, Cairn research) (SI_Figure 1). The camera splitter was equipped with emission filters (ET460/50m, ET525/50m, ET605/70m, and ET700/75m, Chroma Technology) and dichroic mirrors (T495LPXR, T565LPXR and T660LPXR, Chroma Technology) for simultaneous fluorescent recordings of different fluorophores. Cameras were connected to a workstation (Dual Xenon 8-core, 128Gb of memory, a Titan XP video card, and 2 hard drives, 4x NVME 1tb and 8x HDD 8tb) through a Camera Link communication protocol (V3 Firebird Camlink Board). Data acquisition and storage were done with commercial software (NIS-Elements AR 5.11, Nikon Instruments). For transmitted light, a manually controlled quartz halogen lamp was housed in the microscope (FN-LH, Nikon Intruments) (Figure 1D). For epifluorescence measurements, a tunable light source (SPECTRA X Light engine, Lumencor) with excitation filters (395/25, 440/20, 270/24, 510/25, 550/15, 575/25 and 640/30, Chroma Technology) was attached to the microscope head and controlled by computer through the Nis-Elements software. For all the experiments shown in this work, a 25X, 1.1 Numerical aperture (NA), 2 mm working distance (WD) objective was used (CFI75, Apochromat Multi-Photon LWD 25X, Water immersion, Nikon Instruments).

### Freestanding bilayer Microscope (FBM) set-up. Excitation optics (Focus-Laser illumination)

For the focus-laser illumination, three solid state continuous wave lasers (473 nm, 561 nm, and 660 nm GEM lasers, Novanta Photonics) were expanded using pairs of convergent lenses in a Keplerian design (C240TME-A, 8 mm focal length, AC127-025-AB, 25 mm focal length, Thorlabs) (BE in SI_Figure 1), and combined, using an array of dichroic (T495lpxr, T600lpxr, Chroma Technologies), broadband dielectric mirrors (BB111-E01, Thorlabs), and right-angle mirrors (MRA10-E02, Thorlabs) mounted on a custom-made support (BC in SI_Figure 1). The combined beam was fed into a 4 mm galvometer- galvometer scanner (LSKGG4/M, Thorlabs) (Figure 1D, SI_Figure 1). A laser shutter was placed before the GG scan head for fast laser gating (LS6, Uniblitz Electronics). The lasers were operated by SMD12 PSU (Novanta Photonics) controllers connected to the workstation through a PXI serial interface module (PXI-8430/4, National instruments) housed in a PXI chassis (NI PXI-1033, National instruments). The GG scan head was, in turn, operated by a GG controller (Thorlabs) driven by a user-defined signal (DC or sine wave) through a BNC analog output (BNC-2110, National Instruments) also housed in the PXI chassis by an analog output module (PXI-6723, National Instruments). An in-house written software in LabView (National Instruments) was used to control the lasers, the shutter, and the GG. The output of the GG was conjugated to an apodization mask (Annular Mask, Photo Sciences)^54^, which was itself conjugated to the back focal plane of the excitation objective, XO, (TL20X-MPL, 20X, 0.6NA, 5.5 WD, Thorlabs) by relay optics (L1 to L3) using a 4f configuration. L1 (f1=75mm, f2=100mm, Thorlabs), L2 (f1=150mm, f2=100mm, Thorlabs), and L3 (f1=125 mm, f2=125 mm, Thorlabs) were made with 30 mm cage components (Thorlabs) mounted on custom-made adaptor posts for the case of L1 and adaptor plates for L2 and L3. L2 and L3 adaptor plates were, in turn, mounted on a modified breadboard (MB2530/M, Thorlabs) designed to raise and orient the plane of excitation optics orthogonal to the detection optics plane (SI_Figure 1). All custom-made parts, as well as the optomechanical models of the microscope, were designed using computer-assisted design software (Inventor Profesional, AutoDesk). All machined parts were fabricated in Aluminum 6061 by a manufacturing service (Xometry) following our designs.

### Freestanding bilayer Microscope (FBM) set-up. Experimental chamber

The design of the experimental chamber was optimized through many iterations of experimental tests, both imaging and electrophysiology. The chamber (Figure 1B) was 3D printed in-house using a multijet printer (ProJet MJP 3600, 3DSystems) on a proprietary resin (VisiJet M3 Crystal, 3DSystems). The chamber was designed to contain the XO through a conically shaped aperture (Figure 1B and SI_Figure 3A) covered by a silicon rubber film to prevent leakages (High-Purity High-Temperature Silicone Rubber Sheet, 1/32” thickness, 50A Durometer, McMaster-Carr) perforated at the center with a 3 mm punch, to allow the access of the XO. To translate the chamber in space with 1 μm resolution, it was screwed through two M6 screws (Figure 1, SI_Figure 1A and SI_Figure 3A) to a right-angle adaptor plate (3DMS-3X3, Sutter Instruments) mounted on a micromanipulator (MP-285, Sutter Instruments). Parallel to the detection optics plane, two access ports were made to allow for buffer perfusion and pressure regulation (Figure 1) using a hydrostatic manometer made from a modified 30 ml syringe and another micromanipulator (SI_Figure 1). For all the experiments shown in this work, no bottom perfusion was done, and the chamber was connected solely to the manometer. A 3-way valve was added to the tubing connecting the chamber and the manometer to introduce a Ag/AgCl electrode when performing electrophysiological experiments. The top part of the chamber was coupled to the sample holders (Cups), which were also 3D printed in-house by multijet printing (Figure 1B). The mounting was done with vacuum grease (high vacuum grease, Dow Corning) to prevent leaks during electrical measurements. The center of the cup was aligned to a cylindrical hole traversing the entire chamber used for transmitted illumination (SI_Figure 3A). The bottom of the chamber was sealed with a glass coverslip. For the perfusion of solution on top of the cup, two perfusion lines were mounted with wax on the chamber providing access to the cup (Figure 1B). These lines were connected to a custom-made manual perfusion system.

### Electrophysiology set-up

For electrical recordings, the voltage across the lipid bilayer was clamped with an amplifier in whole-cell mode (Axopatch 200B, Axon Instruments) by two Ag/AgCl electrodes placed on the cup and in the manometer line (see previous section). The analog current signal was filtered at 1 kHz (low-pass, Bessel) and digitized at 10 kHz (Digidata 1550B digitizer, Molecular Devices). Digitized data were recorded on a computer using the software pClamp (Molecular Devices) and analyzed using Clampfit (Molecular Devices).

### FEP partition fabrication

FEP sheets (Fluorinated Ethylene Propylene Copolymer - Film, 0.075 mm, Goodfellow) were cut into 1x1 cm squares and indented at the center, either with a weak CO2 laser or a tungsten needle. By sparking the partitions between a high-frequency generator (BD10AS, Electro Technic Products) and a sharp needle, the damaged region on the partition melted, leaving a round and smooth hole (Figure 1A). Changing the duration and intensity of the spark led to hole sizes ranging from 100 to 500 mm. Perforated partitions were sonicated for 30 seconds and stored in ethanol for up to five days.

### Planar bilayers (PBs) formation

Before FBM experiments, a small amount (0.5 μl) of lipid solution was added to both sides of the partition and left to dry for ∼10 minutes. After mounting a cup on the experimental chamber, the entire chamber was filled with imaging buffer (IB, 150 mM KCl, 10 mM Hepes pH 7.4, 2.5 mM Protocatechuic acid, 5 nM Protocatechuic acid dehydrogenase) followed by the addition of the perfusion adaptors. The perfusion lines were purged of air by perfusing 1-2 ml of IB, after which the system was allowed to equilibrate for 5 minutes. Then, FEP partitions were carefully mounted onto the cups with vacuum grease, and the top electrode was inserted into the top chamber, leading to a sharp increase in current. PBs were then made by applying a small amount (1-10 nl) of lipid across the hole of the bilayer. To do this, 1 μl of lipid was first loaded on a pipette and then quickly ejected into the air, leaving a residual lipid solution on the pipette. The pipettes were then inserted into the buffer on the cup, and the bilayer was formed by pushing an air bubble on top of the partition’s hole. The formation of the bilayer was clear from imaging (Figure 1A) and electrical recordings (quick reduction in the current followed by a progressive increase in capacitance). All experiments were done at room temperature. Experiments were done using POPE:POPG (3:1 weight ratio) lipids, except for the phase separation experiments.

### Proteoliposome (PL) fusion

Unless noted otherwise, PLs containing the IMPs of interest were mixed with an equal volume of 1M KCl, briefly sonicated (1 second), added on top of the bilayer, and left to sink for 2 minutes before adding 1 μl of 3M KCl to induce the fusion of vesicles on the bilayer. Unfused PLs were washed away by perfusing 5 ml of IB. For PLs containing G_αi1_ KCl induction was not necessary as we found that G-proteins can diffuse from liposomes into the freestanding bilayer.

### Imaging with the FBM

Focused-laser illumination was turned on after perfusion for hTRAAK and G_αi1_, whilst for M2R and GIRK2, illumination was preceded by the addition of 1 nM labeled NbALFA and another 5 ml of perfusion. The illuminated area was adjusted for every bilayer by changing the input voltage amplitude on the GG. To create a square illumination, two sine waves (10-100 mV) were used, one for the horizontal and one for the vertical scan, at frequencies of 400 and 2500 Hz, respectively (SI_Figure 2C). Videos were recorded, typically at a frame rate of 50 Hz and with a bit depth of 12-bit (12-bit Sensitive mode in Nis-Elements 5.11). Single particle tracking was done using uTrack software to obtain the diffusion coefficient (D) and the anomalous coefficient (α) for every track^55^. Further analysis of the data was done using custom-written software in Matlab (Mathworks). D obtained from the analysis (in pixels^2^/frame) were converted to μm^2^/s using the effective pixel size (0.26 μm for the 25X objective) and the time difference between consecutive frames.

Analysis of the tracks (SI_Figure 4A) to identify the fraction of immobile particles was done by selecting a diffusion cutoff based on the 1% lowest diffusion coefficients (0.5 μm^2^/s) of the distribution for M2R measured with the FBM. Particles with diffusion coefficients (D) lower than this threshold were considered immobile, whilst particles with higher D were considered Brownian. Particles with an anomalous coefficient (α) higher than 1.25 were considered directed. Brownian and confined-Brownian were not distinguished in this analysis. To analyze the single tracks for Figure 2A, videos were cropped around a particle, and single particle tracking was performed using Trackmate (Fiji)^56^. Coordinates were then extracted to calculate the mean squared displacement for different increments of time (MSD vs. τ) and then fitted using equation 1.

### Quartz coverslip preparation

For quartz-supported bilayer experiments, quartz coverslips were plasma-cleaned with oxygen as a processing gas for 0.5-1 minute. Pegylation of Quartz coverslips was done as explained in Chandradoss *et al.*^57^ Briefly, quartz coverslips were cleaned by dipping them in different cleaning solutions, in the order: acetone, 1M KOH, and piranha (H2SO4:H2O2 3:1), followed by washes with deionized water. After cleaning, coverslips were placed in a staining jar with 100 ml of methanol, 5 ml acetic acid, and 3 ml of 3-aminopropyl trimethoxysilane (APTES). After 30 minutes, the solution was replaced by fresh methanol, and this was repeated 3 times. The coverslips were then dried with Nitrogen gas and incubated with a 0.1 M sodium bicarbonate (pH 8.5) solution with 0.6 mM biotinylated NHS-ester PEG (5,000 Da) and 25 mM NHS-ester mPEG (5,000 Da) for 3-5hs. Finally, coverslips were rinsed with deionized water, dried with Nitrogen gas, and stored at -20°C until use. We should note that pegylated quartz coverslips showed fewer fluorescent particles than the other methods, implying that pegylation prevented SB formation to some degree (SI_Video 3-5).

### Mica coverslip preparation

Mica coverslips were prepared by coupling a mica sheet to a quartz coverslip with optical glue as described elsewhere^58^. Briefly, coverslips were cleaned with 2% detergent (Hellmanex III, Hellma Analytics) and ethanol before gluing previously cut mica leaflets (5 mm x 5 mm, 1872-CA, SPI) on top of them with a low viscosity optical adhesive (NOA60, Norland products). After curing the adhesive with one hour exposure to UV light, another coverslip was glued on top of the mica with a high-viscosity optical adhesive (NOA63, Norland products), followed by another round of UV exposure. Coverslips were separated to expose a freshly cleaved mica surface just before performing SB experiments.

### Formation of Supported bilayers (SBs)

To prepare SB from M2R containing liposomes onto the different substrates, we follow a published protocol with a few modifications^34^. Coverslips (prepared as described in the previous sections) were placed inside a 37°C incubator in a pipette box containing wet tissue paper to keep the humidity constant. Proteoliposomes were diluted to 0.1 mg/ml in M2R reconstitution buffer without TCEP before adding 75 μl of the sample onto the coverslips. After 1 minute, an extra 150 μl of buffer was added. After another 2 minutes, the coverslips were washed by adding another 200 ul of buffer, followed by careful mixing, removing, and adding more buffer. This procedure was repeated 3 times.

### SBs TIRF experiments

A separate chamber for TIRF experiments on microscopy coverslips was exchanged for the FBM chamber, keeping the rest of the microscope the same (SI_Figure 3B). The TIRF chamber was designed to hold a Pellin-Broca prism with a 20 mm square aperture (ADB-20, Thorlabs) on top of which the samples (coverslips) were mounted using 10 μl of immersion oil (Type F immersion liquid, Leica Microsystems). After assembling the coverslip, a 3D-printed ring (TIRF - Cup) was secured with vacuum grease and filled with IB for imaging (SI_Figure 3B). Imaging was done as described before for FBM experiments, except that longer exposures were used (120 ms, for mica and Quartz and 130 ms for Pegylated Quartz).

### hTRAAK determination of Po

For Po determination experiments, POPE:POPG (3:1 weight ratio) lipid bilayers were used to match the composition of hTRAAK liposomes. Liposomes were added on top of the bilayers and left to sink and fuse as described before, with the following differences: liposomes were diluted 50 times with IB followed by a 2-fold dilution with 1M KCl to a final concentration of 0.1 mg/ml, and hTRAAK vesicles were allowed to sink for only 30 s before inducing fusion with KCl. This is because hTRAAK-containing liposomes have a stronger proclivity to fuse than all the other proteins we studied. Following the fusion of the PLs, the top chamber was perfused with 5 ml to remove unfused vesicles. Imaging using a scanned-focused laser was then performed as described above. Currents under voltage clamp were measured during the whole procedure. To calculate the number of hTRAAK channels in the bilayer, videos were cropped to a region of the bilayer with homogeneous focus and illumination. Then, uTrack software was used to detect particles within the cropped videos. The number of detections was plotted as a function of time (frames) for each video to estimate the bleaching rate (SI_Figure 7). Then, the average number of particles from different frames was calculated up to a frame that showed less than 5% reduction on the number of particles. The number of detections was divided by the area of the (cropped) frames, which was calculated from the coordinates of the farthest detected particles, to obtain the density of detections. This density was then multiplied by the area of the bilayer, measured using the ROI area measurement feature on the NIS-elements software, and by a correction factor (LR) for the labeling efficiency (15%) to obtain the total number of channels (N) on the bilayer. Potassium currents were measured at 50 mV and divided by N and the single channel current for hTRAAK at that voltage (3.6 pA) to obtain the Po^42^. hTRAAK tension titration experiments were done as described previously without any modification^40^. We note an important assumption in the determination of Po is that incorporated TRAAK channels are functional.

### Phase-separation experiments

4.4 mg of 1,2-diphytanoyl-sn-glycero-3-phosphocholine (DphPC, Avanti Polar Lipids), 2 mg of 1,2-distearoyl-sn-glycero-3-phosphocholine (DSPC, Avanti Polar Lipids) and 1mg of Cholesterol (Co, Avanti Polar Lipids) in chloroform were mixed to a final 2:1:1 molar ratio. At this point, 250 ng of 1,2-distearyl-sn-glycero-3-phosphoethanolamine-N-(TopFluor AF488) (ammonium salt) was added to indicate the ordered phase. The lipid mixture was dried under Argon gas and left under vacuum for at least 2 hours before resuspending into a mixture of Decane:Butanol (9:1) to a final concentration of 22 mg/ml. PBs were formed as described above. After imaging with 2 channels, hTRAAK trajectories were obtained using uTrack software and were overlayed on the first frame of the lipid channel using MatLab (MathWorks). Experiments were done in IB at room temperature.

## Aknowledgements

We thank Anastasios Siskoglou and Daniel C. Gross for assistance and advice in the design and assembly of the FBM, and James Petrillo and Peer Strogies at the Precision Instrumentation Technologies Resource Center for assistance in the fabrication of components for the FBM. Dr. Shiva V. Mandala and Dr. Joël Bloch for useful comments on the manuscript; Dr. Eric Betzig for sharing optomechanical designs, and Dr. Scott Blanchard, Dr. Daniel S. Terry and members of the MacKinnon and Chen labs for useful discussions. R.M. is an investigator of the Howard Hughes Medical Institute. G.P.-M. did part of this work as a Howard Hughes Medical Institute Fellow of the Life Sciences Research Foundation.

## Supporting Figures

**SI_Figure 1.**
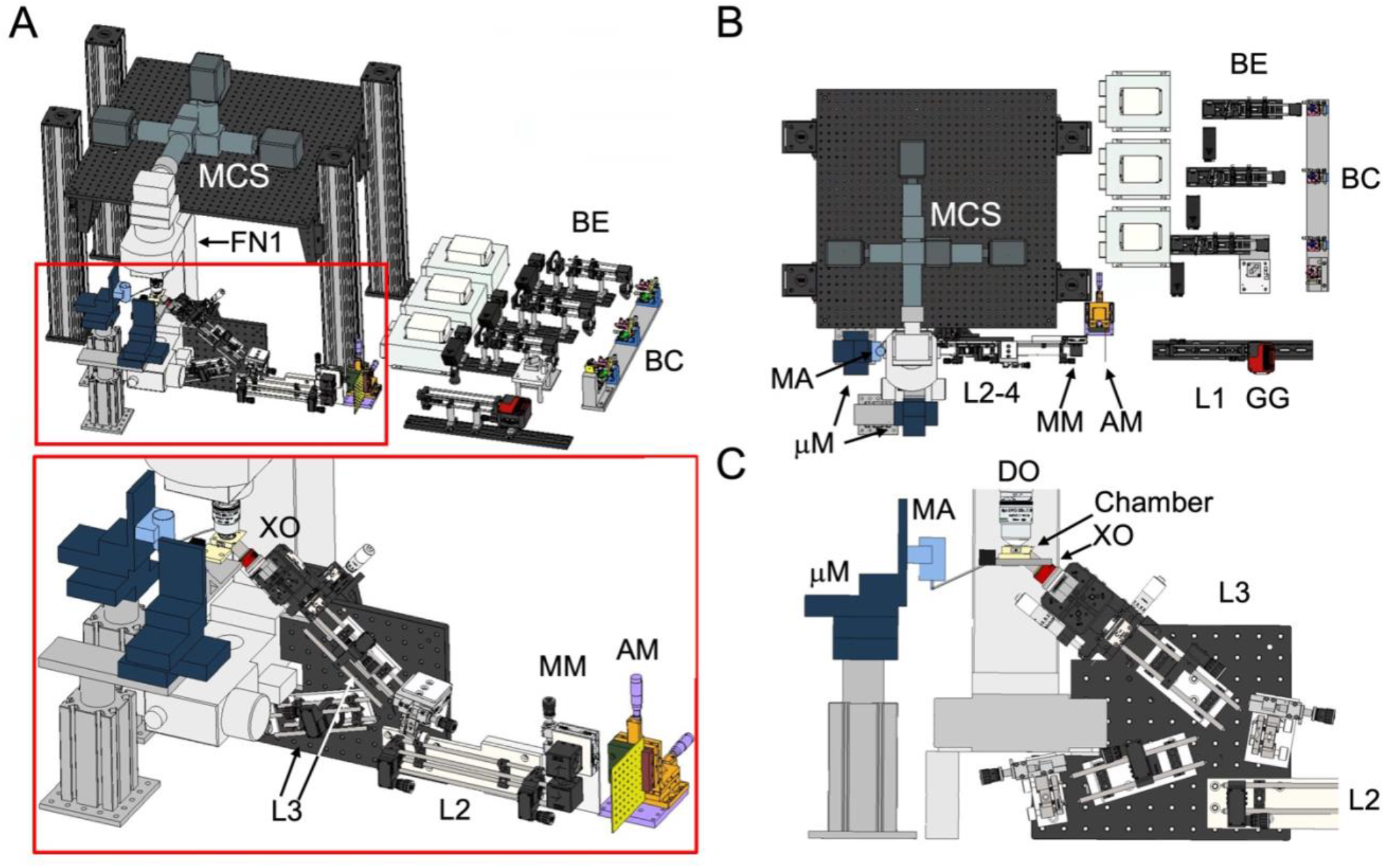
Views of the optomechanical model of the FBM. A. Top-front view. A magnified view of the excitation optics and the experimental chambers is shown in the red box. **B.** Top view. **C.** Cross-sectional view of the excitation optics and experimental chamber. Abbreviations stand for BE: Beam expanders, BC: Beam (laser) combiner, GG: Galvo-Galvo scanner, L1-L3: Optical relays, AM: Apodization mask, MM: movable mirror, mM: micromanipulator, MA: manometer, MCS: multi-camera splitter, XO: excitation objective, DO: detection objective, FN1: Upright microscope from Nikon (model FN1).

**SI_Figure 2.**
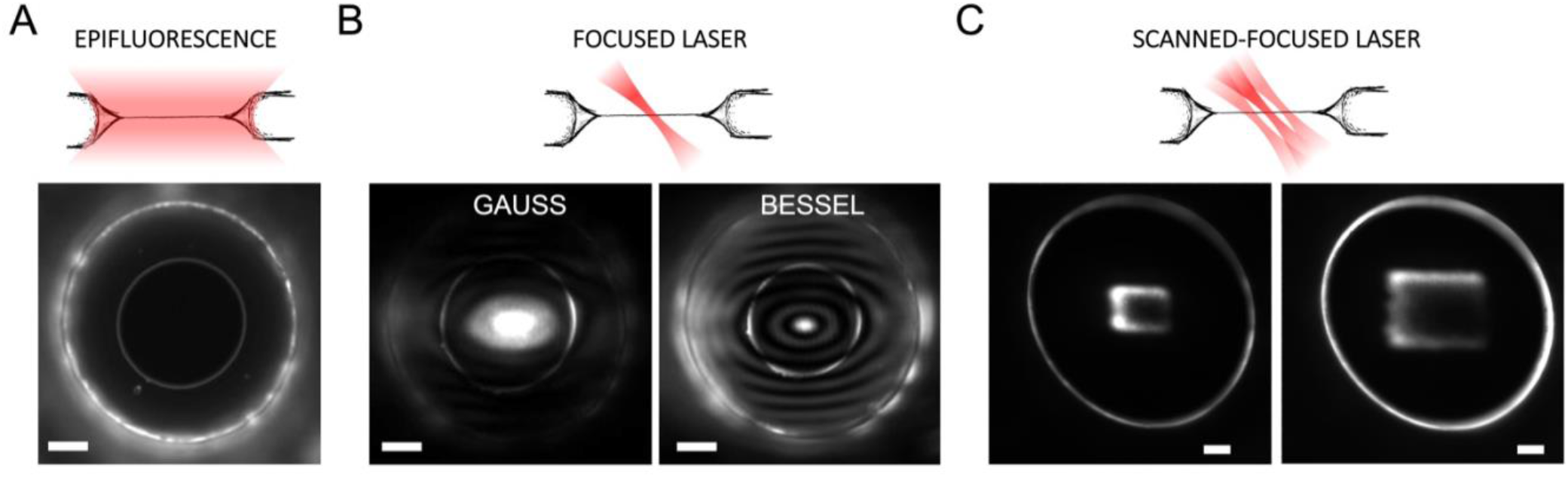
Comparison of different modes of illumination of a POPE: POPG (3:1 by weight) bilayer with Rhodamine-DOPE (0.01% w/w). A. Epifluorescence illumination. **B.** Focused-laser illumination with a Gauss (left) and Bessel (right) beam. **C.** Scanned focused-laser illumination using a scanning amplitude of 50 (left) and 100 (right) mV. Images A and B were taken on the same bilayer, and C was on a different one. Scale bars: 10 μm.

**SI_Figure 3.**
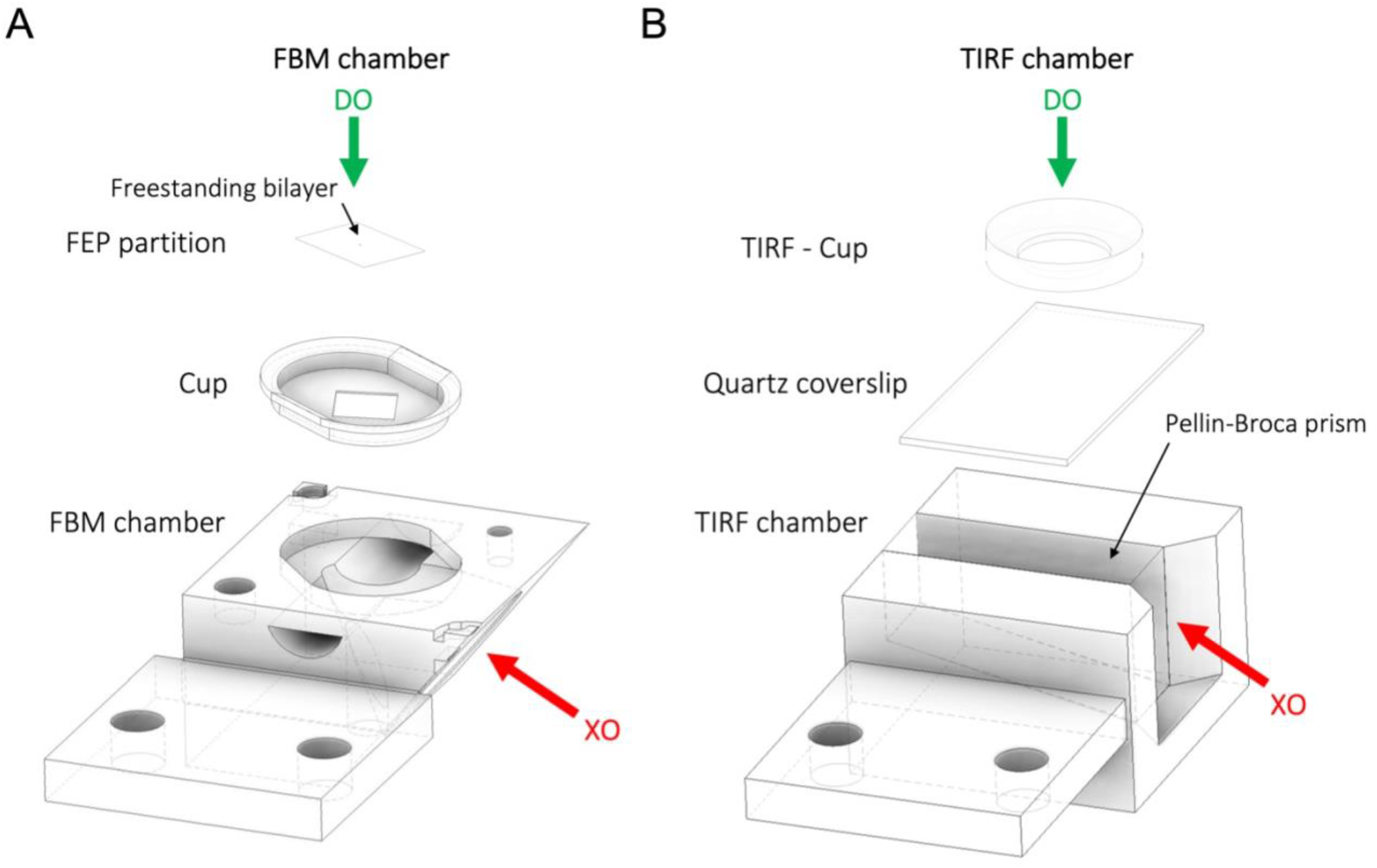
Optomechanical models of FBM and TIRF chambers. A. FBM chamber. Amplified views of the FBM chamber depicting the exchangeable cup that separates the top and bottom chamber and the FEP partition where the freestanding bilayer is formed. **B. TIRF chamber.** Amplified view of the TIRF chamber used for the SB experiments depicting the quartz coverslip on which SBs were formed and the TIRF cup that contains the imaging buffer. The coverslip is coupled to a Pellin-Broca prism through immersion oil. Red and green arrows are used to represent the directions of the incoming excitation (XO) and detection optics (DO), respectively.

**SI_Figure 4.**
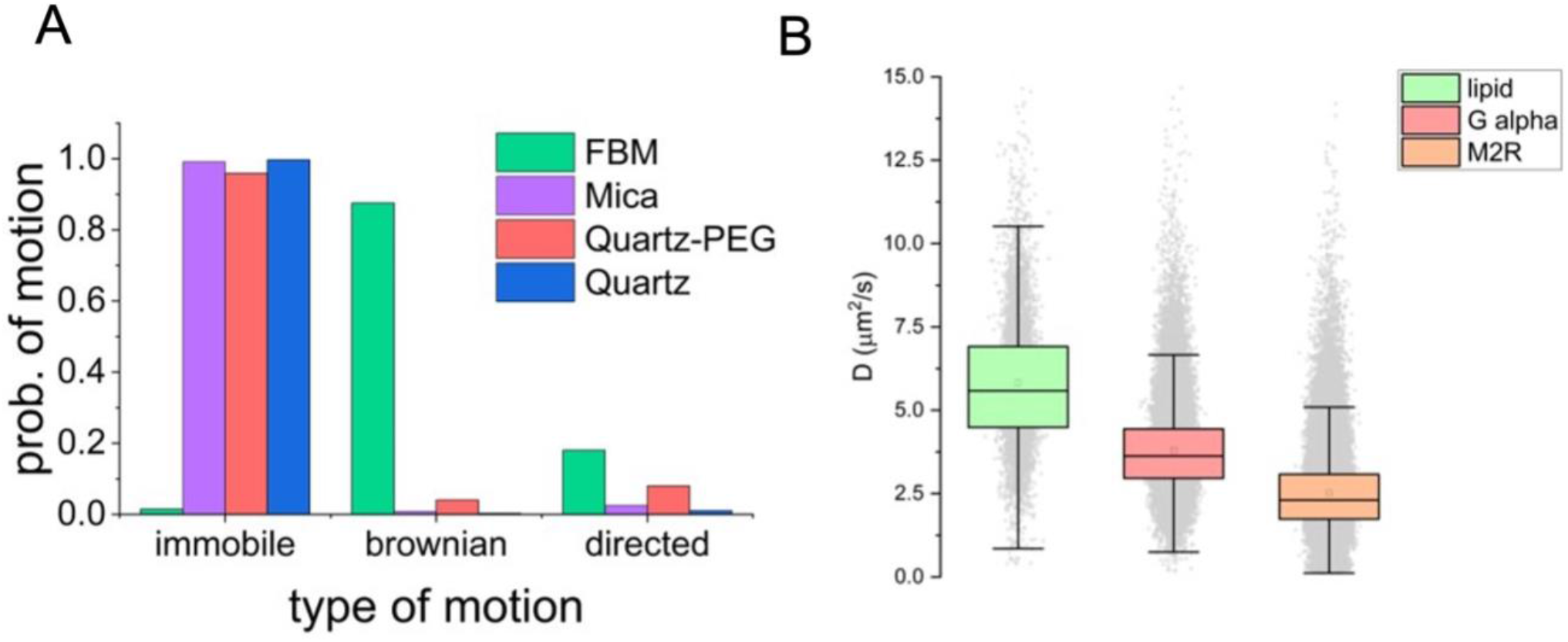
Single-particle-tracking (SPT) experiments. A. Classification of the data on M2R shown in. figure 2B. Results show a significant difference between the dominant type of motion in the FBM (Brownian) and SBs (immobile). **B. Diffusion coefficients (D) for DOPE-Cy5 (green), G_αi_ (red) and M2R (orange) in the FBM**. The mean, 25;75 percentile (Box) and 5;95 percentile (bars) are shown overlaying the data. G_αi_ and M_2_R were labeled with LD655 through Sfp site-directed labeling and with NbALFA-LD655, respectively.

**SI_Figure 5.**
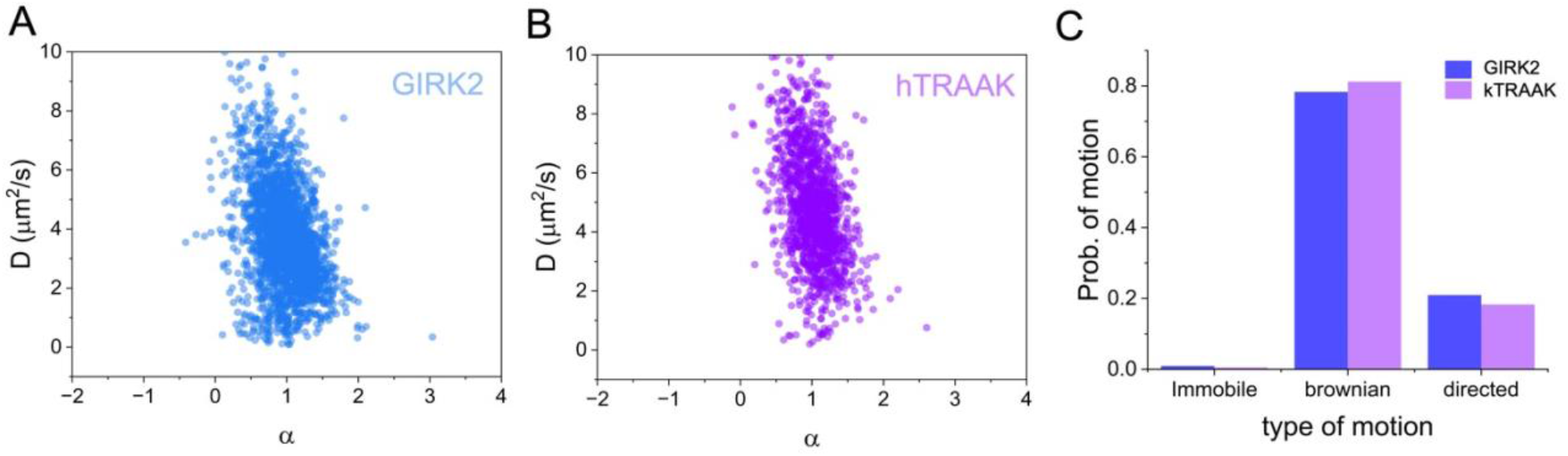
Single-particle-tracking (SPT) experiments. A. Diffusion coefficients (D) vs. anomalous coefficient (α) for GIRK2 (A) and hTRAAK (B). C. Classification of the tracks shown in panels A **and B for GIRK2 (blue) and hTRAAK (violet).** The results indicate that both proteins remain mobile in FBM experiments. GIRK2 and hTRAAK where labeled with NbALFA-LD655 and LD655-NHS, respectively.

**SI_Figure 6.**
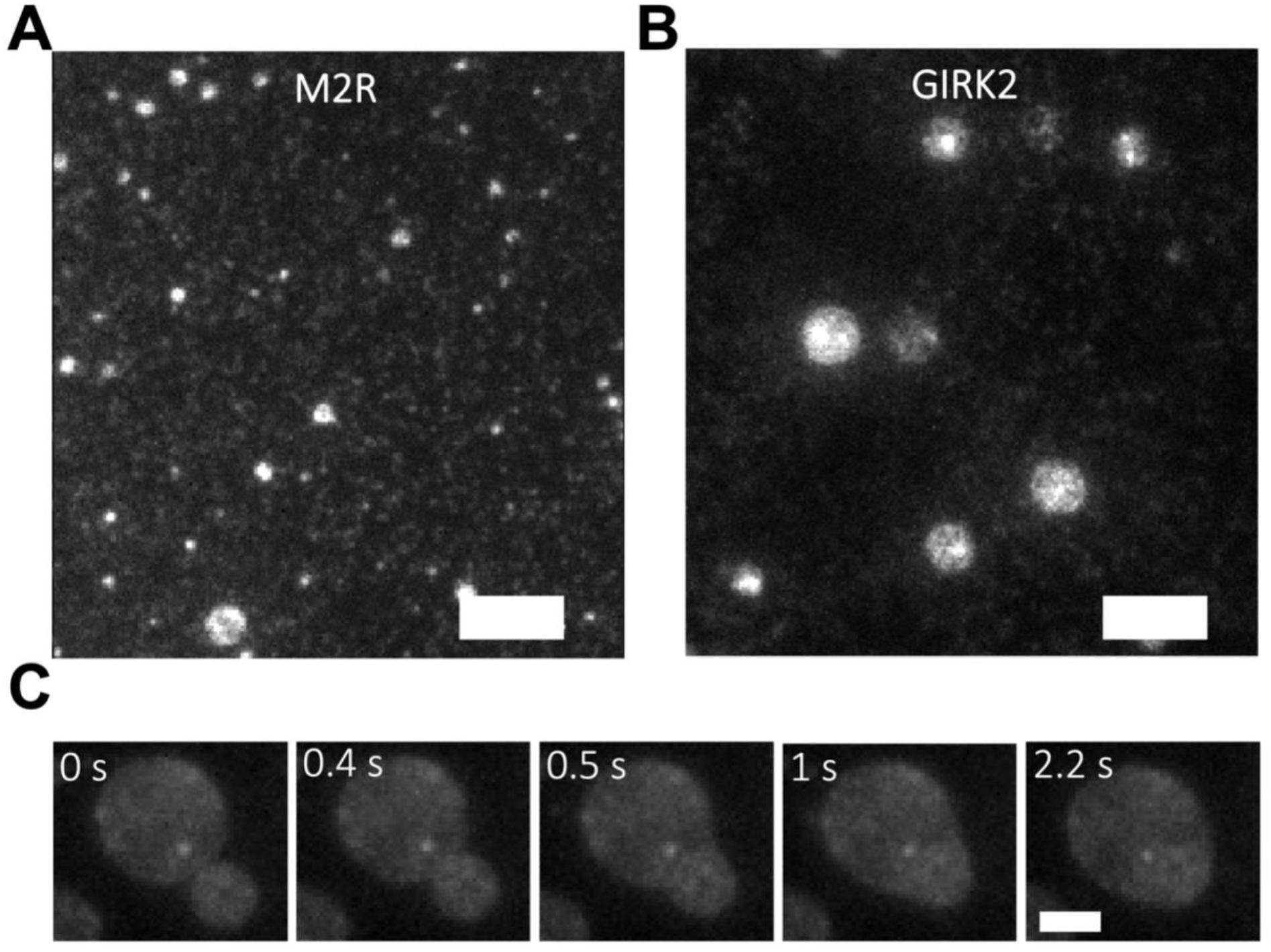
Planar protein aggregates are observed in FBs after proteoliposome fusion. Aggregates of 1-10 μm are routinely observed after vesicle fusion for **M2R (A)** and **GIRK2** (**B). C. A time sequence showing the fusion of two planar aggregates of GIRK2.** Scale bars: 10 μm (A, B) 4 μm (C).

**SI_Figure 7.**
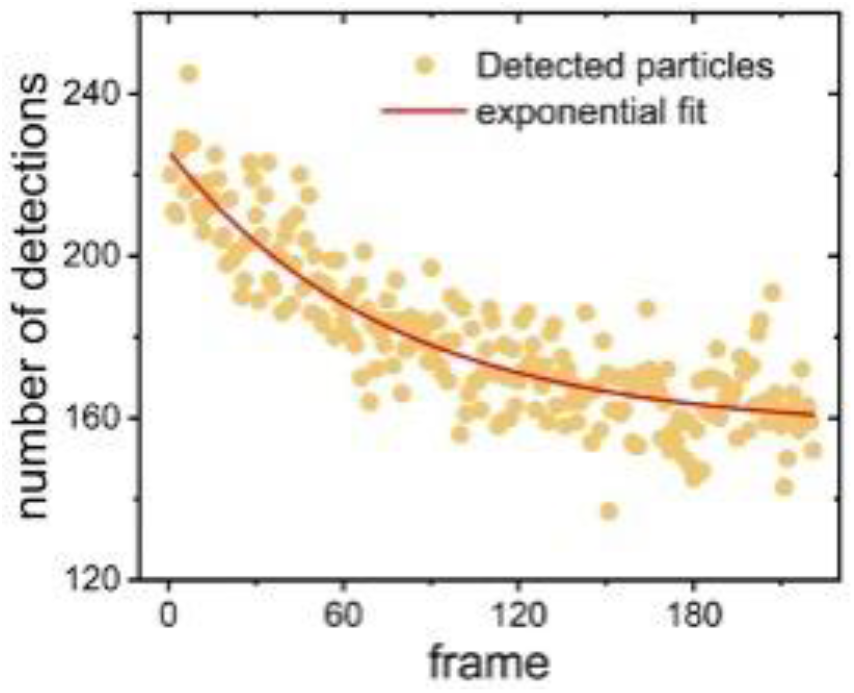
**Number of hTRAAK particles detected per frame during a FBM video**. Each frame lasts 30 ms. An exponential fit to the data is shown as a red line (r^2^=0.99).

## Supporting information

Videos were converted and compressed from 12 bit (.nd2 format) to 8 bit (.mp4 format) to ease their visualization with common video players.

**Supporting video 1.** Freestanding bilayer containing LD655-labeled hTRAAK observed under epifluorescence. Frame rate: 50 Hz.

**Supporting video 2.** Freestanding bilayer containing LD655-labeled hTRAAK observed under focused-laser illumination. Frame rate: 50 Hz.

**Supporting video 3.** Quartz-supported bilayer observed under TIRF illumination. Frame rate: 8 Hz. **Supporting video 4.** Mica-supported bilayer observed under TIRF illumination. Frame rate: 8 Hz. **Supporting video 5.** Pegylated-Quartz-supported bilayer observed under TIRF illumination. Frame rate: 8 Hz.

**Supporting video 6.** Freestanding bilayer containing the G-protein coupled receptor M2R labeled with NbALFA-LD655. Frame rate: 67 Hz.

**Supporting video 7.** Freestanding bilayer containing the G-protein alpha G_αi1_ labeled with LD655 through Sfp-mediated site-directed labeling. Frame rate: 68 Hz.

**Supporting video 8.** Freestanding bilayer containing the fluorescent lipid DOPE-Cy5. Frame rate: 50 Hz.

